# Ubiquity and origins of structural maintenance of chromosomes (SMC) proteins in eukaryotes

**DOI:** 10.1101/2021.05.15.444277

**Authors:** Mari Yoshinaga, Yuji Inagaki

## Abstract

Structural maintenance of chromosomes (SMC) protein complexes are common in Bacteria, Archaea, and Eukaryota. SMC proteins, together with the proteins related to SMC (SMC-related proteins), constitute a superfamily of ATPases. Bacteria/Archaea and Eukaryotes are distinctive from one another in terms of the repertory of SMC proteins. A single type of SMC protein is dimerized in the bacterial and archaeal complexes, whereas eukaryotes possess six distinct SMC subfamilies (SMC1-6), constituting three heterodimeric complexes, namely cohesin, condensin, and SMC5/6 complex. Thus, to bridge the homodimeric SMC complexes in Bacteria and Archaea to the heterodimeric SMC complexes in Eukaryota, we need to invoke multiple duplications of an SMC gene followed by functional divergence. However, to our knowledge, the evolution of the SMC proteins in Eukaryota had not been examined for more than a decade. In this study, we reexamined the ubiquity of SMC1-6 in phylogenetically diverse eukaryotes that cover the major eukaryotic taxonomic groups recognized to date and provide two novel insights into the SMC evolution in eukaryotes. First, multiple secondary losses of SMC5 and SMC6 occurred in the eukaryotic evolution. Second, the SMC proteins constituting cohesin and condensin (i.e., SMC1-4), and SMC5 and SMC6 were derived from closely related but distinct ancestral proteins. Based on the above-mentioned findings, we discuss how SMC1-6 have diverged from the archaeal homologs.

## INTRODUCTION

Chromosomes comprise DNA molecules, which are the body of genetic information, and a large number of proteins with diverse functions. In eukaryotes, cohesin and condensin, together with many other proteins, maintain the integrity of chromosome structure. Cohesin and condensin participate in protein complexes (Anderson et al. 2002) that bundle sister chromosomes together during mitosis (Haering et al. 2008) and meiosis (Ishiguro 2019), and aggregate chromosomes (Sutani and Yanagida 1997), respectively. Cohesin is constituted by two Structural Maintenance of Chromosomes (SMC) proteins (SMC1 and SMC3) (Losada et al. 1998) and accessory subunits Rad21/Scc1 and STAG1/Scc3 (Birkenbihl and Subramani 1995; Carramolino et al. 1997; Tóth et al. 1999). Condensin contains SMC2 and SMC4 (Hirano and Mitchison 1994), and a different set of accessory subunits CAP-D2, CAP-G, and CAP-H (Hirano et al. 1997). There are two additional SMC proteins, SMC5 and SMC6 (Lehmann et al. 1995; Fousteri and Lehmann 2000), which comprise the “SMC5/6” complex together with six accessory proteins (Nse1-6) (Andrews et al. 2005; Fujioka et al. 2002; Hu et al. 2005; Pebernard et al. 2004; Pebernard et al. 2006) and involve mainly in DNA repair but also replication fork stability (Aragón 2018).

SMC proteins, together with MukB, Rad50, and RecN, belong to a large ATPase superfamily with unique structural characteristics (Niki et al. 1991; Funayama et al. 1999; Löwe et al. 2001). SMC proteins comprise “head” that hydrolyzes ATP, “hinge” that facilitates the dimerization of two SMC proteins (SMC1 and SMC3 in cohesin, SMC2 and SMC4 in condensin, and SMC5 and SMC6 in the SMC5/6 complex), and antiparallel coiled coils connecting the head and hinge (Melby et al. 1998). As ATPases, SMC proteins bear eight motifs such as Walker A (P-loop), Walker B, ABC signature motif (C-loop), A-loop, D-loop, H-loop (switch motif), R-loop, and Q-loop, all of which are required for ATP binding and hydrolysis. In the ATPases belonging to the SMC superfamily, the Walker A motif, A-loop, R-loop, and Q-loop are located at the N-terminus of the molecule, being remote from the rest of the motifs at the C-terminus (Palou et al. 2018). Thus, SMC proteins most likely form hairpin-like structures to make all of the sequence motifs for ATP binding in close proximity in the tertiary structures (Melby et al. 1998).

The vast majority of the members of Bacteria and Archaea possess a single SMC protein for DNA strand aggregation. In contrast to the eukaryotic SMC complexes containing heterodimeric SMC proteins, the SMC complexes in Bacteria and Archaea comprise two identical SMC proteins (i.e., homodimeric), together with accessory subunits (Britton et al. 1998; Soppa 2001). It is noteworthy that the SMC protein is not conserved strictly in Bacteria or Archaea (Soppa 2001). For instance, the absence of the conventional SMC protein in the Crenarchaeota genus *Sulfolobus* was experimentally shown to be complemented by the proteins that are distantly related to the authentic SMC, namely coalescin (Takemata et al. 2019).

To our knowledge, no study on the diversity and evolution of SMC proteins sampled from phylogenetically diverse eukaryotes has been done since Cobbe and Heck (2004). Their phylogenetic analyses recovered individual clades of SMC1-6, and further united (i) SMC1 and SMC4 clades, (ii) SMC2 and SMC3 clades, and (iii) SMC5 and SMC6 clades together. Henceforth here in this work, we designated the three unions as “SMC1+4 clan,” “SMC2+3 clan,” and “SMC5+6 clan,” respectively. Based on the phylogenies inferred from the SMC proteins in the three domains of Life, the authors proposed that SMC1-6 were yielded through gene duplication events that occurred in the early eukaryotic evolution.

The pioneering work by Cobbe and Heck (2004) was a significant first step to decipher the origin and evolution of the six SMC subfamilies in eukaryotes, albeit they provided no clear scenario explaining how a primordial SMC protein diversified into SMC1-6 prior to the divergence of the extant eukaryotes. Thus, we reassessed the ubiquity and the phylogenetic relationship among the six eukaryotic SMC subfamilies in this study. Fortunately, recent advances in sequencing technology allow us to search for SMC homologs in the transcriptome and/or genome data of phylogenetically much broader eukaryotic lineages than those sampled from metazoans, fungi (including a microsporidian), land plants, and trypanosomatids analyzed in Cobbe and Heck (2004). Furthermore, computer programs for the maximum-likelihood (ML) phylogenetic methods, as well as hardware, have been improved significantly since 2004. Thus, hundreds of SMC proteins from diverse eukaryotes can be subjected to the ML analyses now, in contrast to Cobbe and Heck (2004) wherein only a distance tree was inferred from the alignment of 148 SMC sequences.

Our survey of SMC1-6 in 101 eukaryotes confirmed the early divergence of the six SMC subfamilies in eukaryotes, albeit the secondary loss of SMC5 and SMC6 most likely has occurred in separate branches of the tree of eukaryotes. Moreover, the phylogenetic analysis of SMC1-6, bacterial and archaeal SMC, and Rad50/SbcC (397 sequences in total) disfavored the single origin of the six SMC subfamilies in eukaryotes and instead suggested that the ancestral molecule of SMC5 and SMC6 is distinct from that of SMC1-4. We finally proposed a scenario for the evolution of SMC in Archaea and Eukaryota.

## RESULTS

We first reexamined the conservation of the six SMC subfamilies in 59 phylogenetically diverse eukaryotes. Among the 59 eukaryotes, we retrieved 366 SMC proteins in total after eliminating non-SMC sequences by preliminary phylogenetic analysis. Subsequently, we classified the 366 SMC proteins into individual subfamilies (i.e., SMC1-6) based on the phylogenetic affinity to the SMC1-6 proteins known prior to this study (Table S1). Table S1 contains truncated SMC sequences likely due to (i) the absence of the full-length transcript in the RNA-seq data and/or (ii) an intron (or introns) that hinders recovery of the entire SMC-coding region from the genome data.

### Inventories of the proteins constituting cohesin and condensin

At least two of SMC1-4 were detected in 52 out of the 59 eukaryotes examined here (Fig. 1: The 7 eukaryotes, from which the set of SMC1-4 failed to be completed, are described in the latter part of this section). A preliminary phylogenetic analysis suggested that lineage-specific gene duplications produced two different types of SMC4 in the ciliate *Oxytricha trifallax* and the parabasalid *Trichomonas vaginalis* (EJY74552.1 and EJY65570; XP_001328834.1 and XP_001328078.1), and those of SMC2 in the ciliate *Paramecium tetrauelia* (XP 001443869.1 and XP 001447973). Likewise, the duplication of SMC6 gene may have occurred independently in two land plants *Arabidopsis lyrata* and *Physcomitrella patens*, yielding two distinctive types of SMC6 (XP 020877425.1 and XP 020884092.1; XP 024388470.1 and XP_024388471.1).

**Figure 1.**
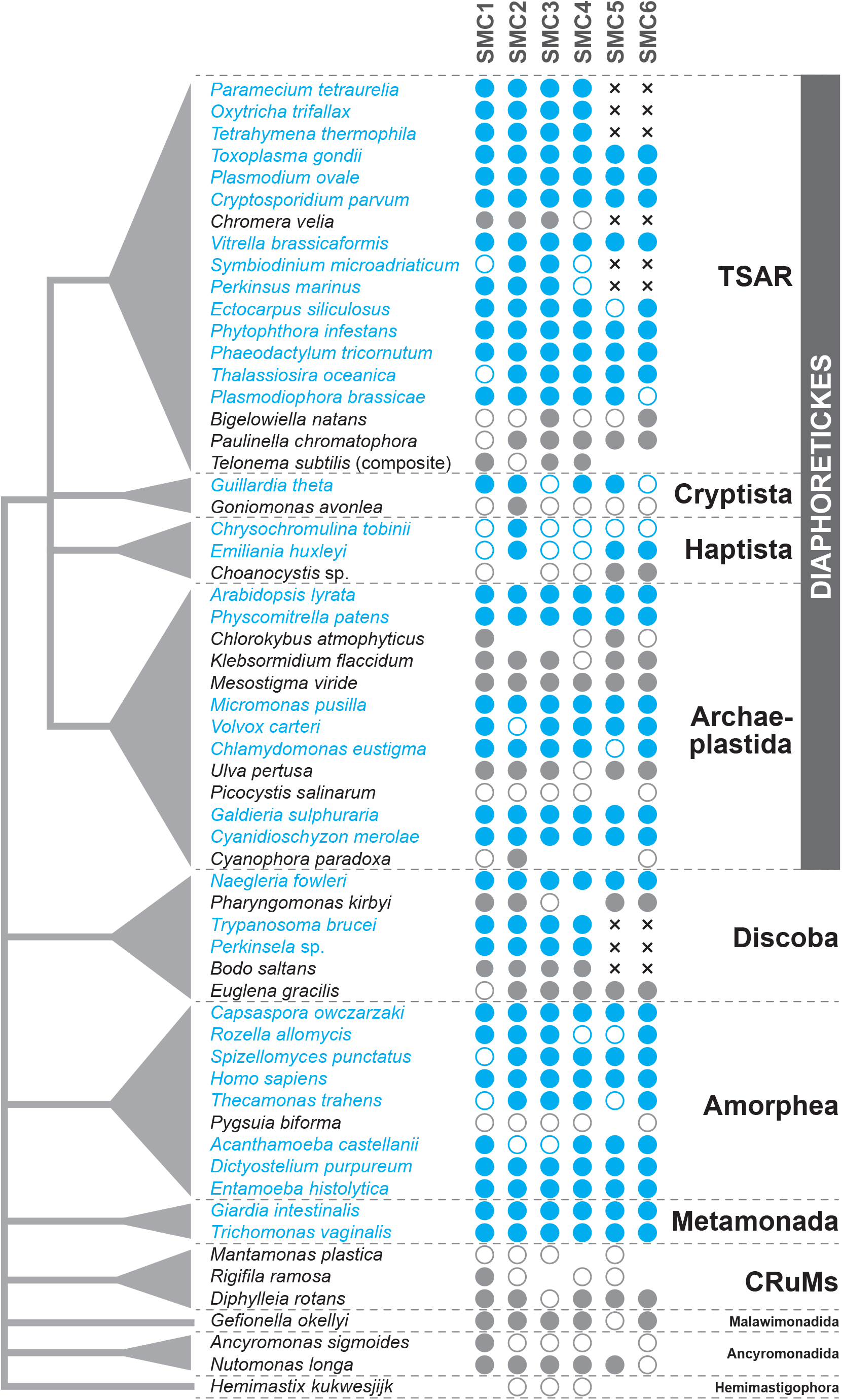
Inventories of the six SMC subfamilies in eukaryotes. The SMC sequences were searched for in 59 species that represent the major taxonomic assemblages in the tree of eukaryotes. The phylogenetic relationship among the major taxonomic assemblages (left) is unsettled except the node uniting TSAR, Cryptista, Haptista, and Archaeplastida, which are so-called Diaphoretickes as a whole. On the right, the presence/absence of the six SMC subfamilies is shown by symbols for each species. Blue (filled) circles represent the full-length SMC amino acid sequences retrieved from the GenBank nr database. Grey (filled) circles represent that the transcripts encoding the full-length proteins were found in the corresponding transcriptome data deposited in the SRA database in the GenBank. Open circles with blue borders represent truncated SMC amino acid sequences found in the GenBank nr database, while those with grey borders represent the truncated transcripts encoding SMC proteins identified in the transcriptome data. We only propose the absence of SMC5 and SMC6 in 9 eukaryotes (highlighted by x-marks), as neither of the two SMC subfamilies was found in their genome or transcriptome data. In case of a certain subfamily being undetected in the transcriptome data, we left its presence/absence uncertain and gave no symbol in the corresponding column in the figure.

From each of the telonemid *Telonema subtilis* and the ancyromonad *Ancyromonas sigmoides*, we detected two separate transcripts, one encoding the N-terminal portion of SMC1 and the other encoding the C-terminal portion of the same protein. Although we found no transcript encoding the entire SMC1, we regard that both *Telonema subtilis* and *Ancryromonas sigmoides* possess the standard SMC1 with the ATP binding motif split into the N and C-termini of a single molecule. Likewise, for SMC1 and SMC3 of the chrompodellid *Chromera velia*, SMC2 of the glaucophyte *Cyanophora paradoxa*, and SMC6 of the heterolobosean *Naegleria fowleri*, we failed to retrieve any transcript/contig encoding the entire protein in the transcriptome/genome data. Although the uncertainties described above could not be resolved at this point, we considered that the SMC proteins mentioned above are of full-length.

We detected a subset of SMC1-4 in *Chlorokybus atmophyticus* (a green alga, Archaeplastida), *Cyanophora paradoxa*, (a glaucophyte, Archaeplastida), *Choanocystis* sp. (a centrohelid, Haptista), *Pharyngomonas kirbyi* (a heterolobosean, Discoba), *Rigifila ramosa* and *Mantamonas plastica* in the CRuMs clade, and the hemimastigophoran *Hemimastix kukwesjijk* (Fig. 1). It is most likely that the 7 species described above do possess SMC1-4 but subsets of subfamilies in them were escaped from our survey due to some experimental reasons. The four SMC subfamilies were found in the phylogenetic relatives of the species mentioned above, except for *Hemimaxtix kukwesjijk* that showed no clear phylogenetic affinity to any other eukaryotes known to date (Lax et al. 2018).

For instance, *Pharyngomonas kirbyi* is closely related to *Naegleria fowleri*, both of which belong to Heterolobosea, and SMC4 was not detected in the former while all of SMC1-4 were completed in the latter (Fig. 1). One of the most plausible reasons for missing one or two out of SMC1-4 in this study is the quality and quantity of sequence data. We suspect that, depending on data quality and/or quantity, all of the SMC subfamilies were difficult to detect in the species for which only transcriptome data were available. In sum, despite the uncertainties above, we anticipate that all eukaryotes have the set of SMC1-4, reinforcing the significance of cohesin (including SMC1 and 3) and condensin (including SMC2 and 4) for cellular viability. The ubiquity of SMC1-4 strongly suggests that SMC1-4 were present in the last eukaryotic common ancestor (LECA).

We also investigated the conservation of accessory subunits of cohesin and condensin in the transcriptome and/or genomic data from the 59 eukaryotes (Fig. S1). In cohesin, SMC1 and SMC3 interact with Rad21/Scc1 and STAG1/Scc3. We detected Rad21/Scc1 and STAG1/Scc3 homologs in 44 and 43 out of the 59 eukaryotes investigated here, respectively. It is worthy to note that the two accessory subunits were detected rarely in the members of Alveolata. This result implies that Rad21/Scc1 and/or STAG1/Scc3 in alveolates are highly diverged and are difficult to detect based on amino acid sequence similarity.

Alternatively, some of the eukaryotes (e.g., alveolates) might have lost Rad21/Scc1 and/or STAG1/Scc3 or substitute the two proteins with as-yet-unknown proteins. Condensin comprises CAP-D2, CAP-G, and CAP-H, together with SMC2 and SMC4. CAP-D2, CAP-G, and CAP-H were found in 48, 33, and 42 out of the 59 eukaryotes, respectively (Fig. S1). Curiously, we failed to find CAP-G in any of the members of Alveolata examined here. Likewise, neither CAP-G nor CAP-H was found in the members of Kinetoplastea examined here. Future experimental studies are necessary to examine the requirement of CAP-G in alveolates, and CAP-G and CAP-H in kinetoplastids. We conducted the phylogenetic analysis of the kleisin superfamily, which is composed of Rad21/Scc1, CAP-H, Nse4, and ScpA in cohesin, condensin, the SMC5/6 complex, and the bacterial/archaeal SMC complex, respectively (Schleiffer et al. 2003). Rad21/Scc1 and CAP-H sequences formed distinct clades, which were sister to each other, in the kleisin phylogeny (Fig. S2).

### Inventories of the proteins constituting the SMC5/6 complex

SMC5 and/or SMC6 were detected in 49 out of the 59 species assessed in this study (Fig. 1). A set of SMC5 and SMC6 was found in 43 out of the 49 species, while either of the two SMC proteins is missing in 6 species (Fig. 1). We suspected that one of the two proteins escaped from our survey in the 6 species because both SMC5 and SMC6 are indispensable to constitute the SMC5/6 complex. Despite the exceptions described above, both SMC5 and SMC6 were found in phylogenetically broad eukaryotes, suggesting that the two SMC subfamilies were present in the LECA.

Neither SMC5 nor SMC6 was detected from 11 out of the 59 eukaryotes examined here (Fig. 1). Curiously, both high-quality transcriptome and genome data are available for 9 out of the 11 eukaryotes, namely three ciliates (*Paramecium tetraurelia, Oxytricha trifallax*, and *Tetrahymena thermophila*), and three kinetoplastids (*Trypanosoma brucei, Bodo saltans*, and *Perkinsela* sp.), the chrompodellid *Chromera velia*, the dinoflagellate *Symbiodinium microadriaticum*, and *Perkinsus marinus* that represents Perkinosozoa, a taxonomic group basal to dinoflagellates (Fig. 1). The failure in identifying SMC5 or SMC6 in the three kinetoplastids is consistent with Gluenz et al. (2008) which detected neither SMC5 nor SMC6 in *Trypanosoma brucei*. We here regard the aforementioned eukaryotes as the candidate lineages that discarded SMC5 and SMC6 secondarily. To pursue the possibility described above, we expanded the search on the repertoire of SMC subfamilies in additional ciliates, dinoflagellates, and chrompodellids, and kinetoplastids and their close relatives (see below). The results of the aforementioned surveys of the SMC subfamilies in the lineages are described in the next section. Unfortunately, we cannot be sure whether both SMC5 and SMC6 are absent truly in *Hemimaxtix kukwesjijk* and *Telonema subtilis* based on the survey solely in the transcriptome data.

The SMC5/6 complex is known to be associated with Nse1–6. In this study, we searched for the Nse1-4 sequences in the 59 eukaryotes, albeit Nse5 and Nse6, both of which are little conserved among eukaryotes (Diaz and Pecinka 2018), were excluded from the survey. The survey of Nse1-4 was less successful than those of the accessory subunits in cohesin and condensin (see above). We found the set of the four Nse proteins only in 8 out of the 59 eukaryotes examined (Fig. S1). Nevertheless, at least one of Nse1-4 was found in phylogenetically diverse eukaryotes, implying that the LECA possessed the SMC5/6 complex associated with at least Nse1-4. The phylogenetic analysis of the proteins belonging to the kleisin superfamily recovered the clade of Nse4 sequences, which was further connected to the Rad21/Scc1 and CAP-H clades excluding the ScpA sequences (Fig. S2). Nse1 and Nse3 are known to bear the similarity to ScpB (an accessory subunit of bacterial/archaeal SMC complex) at the tertiary structural level (Palecek and Gruber 2015). However, it was difficult to detect the similarity among Nse1, Nse3, and ScpB at the amino acid sequence level and we could not assess their phylogenetic relationship.

### Secondary losses of SMC5 and SMC6

The survey of the SMC subfamilies illuminated putative secondary losses of SMC5/6 in kinetoplastids, ciliates, dinoflagellates plus *Perkinsus*, and a chrompodellid *Chromera velia* (Fig. 1). Thus, we additionally surveyed the SMC subfamilies in the transcriptome data of three kinetoplastids plus three of their relatives (two diplonemids and a euglenid), 15 ciliates, three chrompodellids, and 18 dinoflagellates. The transcriptome data of the species additionally surveyed were retrieved from public databases and assembled into the contig data. As a transcriptome analysis unlikely covers the entire repertory of the genes in the corresponding genome, all of the SMC subfamilies in the species of interest are not necessarily detectable by the survey in the contig data. Nevertheless, if SMC5/6 is truly absent in a particular group of eukaryotes, neither SMC5 nor SMC6 is unlikely detected in any members of the group of interest.

We examined whether the putative absence of SMC5/6 in kinetoplastids *Trypanosoma brucei, Perkinsela* sp., and *Bodo saltans* (Fig. 1) is applicable to other members of the class Kinetoplastea and members belonging to classes Diplonemea and Euglenida that comprise the phylum Euglenozoa with Kinetoplastea. As anticipated, in all of the kinetoplastids and diplonemids additionally examined, SMC5 or SMC6 was not detected while SMC1-4 were found (Fig. 2A). In contrast, SMC1-6 were detected in the sequence data from two euglenids *Euglena gracilis* and *Eutreptiella gymnastics* (Fig. 2A). In the tree of Euglenozoa, Kinetoplastea and Diplonemea branch together by excluding Euglenida. Combining the presence/absence of SMC5/6 and the phylogenetic relationship among the three classes in Euglenozoa together, we propose that a single loss of SMC5/6 occurred in the common ancestor of kinetoplastids and diplonemids.

**Figure 2.**
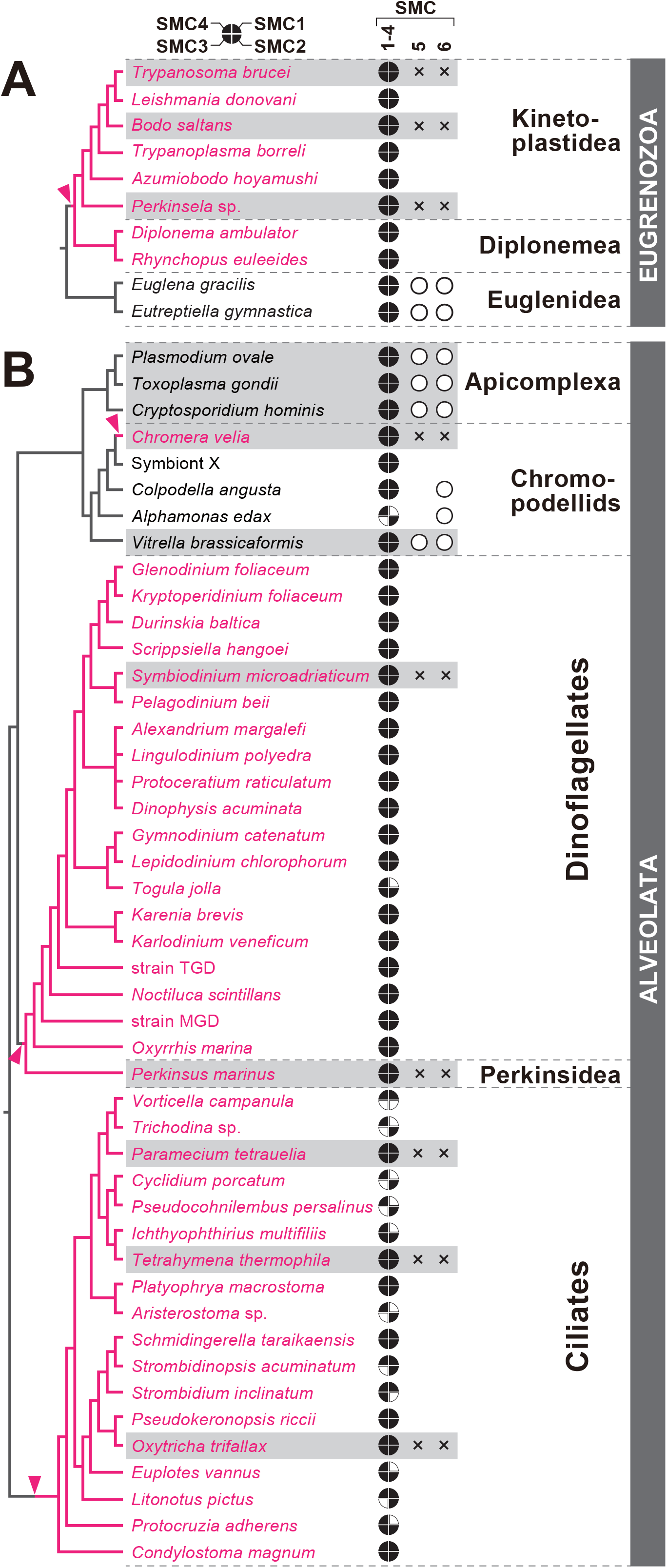
Taxonomic groups lacking SMC5 and SMC6. For each species examined here, the presence of SMC1-4 is represented by filled portions of the circle (upper right, SMC1; lower right, SMC2; lower left, SMC3; upper left, SMC4), regardless of whether they are full-length or truncated. If a subset of SMC1-4 was not detected in the transcriptome data, the corresponding portion(s) is left blank. Open circles indicate that we detected the full-length or truncated transcripts encoding SMC5/SMC6. According to the rule described in the legend for Figure 1, we judged the absence of SMC5 and/or SMC6 (highlighted by x-marks). In case of no transcript encoding SMC5/SMC6 being detected in a certain transcriptome, the corresponding column is left blank. The species, for which both genome and transcriptome data are available, are shaded in grey. The secondary losses of SMC5 and SMC6 were predicted on the branches highlighted by arrowheads. The descendants of the organisms, which discarded SMC5 and SMC6 secondarily, are colored in magenta. (A) Inventories of SMC1-6 in the members of Euglenozoa. The phylogenetic relationship among the kinetoplastids considered here is based on Yazaki et al. (2017). (B) Inventories of SMC1-6 in Alveolata. The phylogenetic relationship among the apicomplexans and chrompodellids, that among dinoflagellates, and that among ciliates are based on Janouškovec et al. (2019), Sarai et al. (2020), and Gao et al. (2016), respectively.

The results from the survey of SMC subfamilies in dinoflagellates were straightforward. Neither SMC5 nor SMC6 was found in any of the dinoflagellate species examined, whereas SMC1-4 were completed in all of them except *Togula jolla*, from which SMC1 was not found (Fig. 2B). Thus, we interpret the result as both SMC5 and SMC6 being lost in the common ancestor of dinoflagellates and perkinsids.

The presence/absence of SMC5 and SMC6 in chrompodellids is likely complicated. Importantly, both SMC5 and SMC6 were identified in *Vitrella brassicaformis* (Fig. 1). Among the species additionally examined, SMC6 was detected in *Alphamonas edax* and *Colpodella angusta* (Fig. 2B). Thus, the ancestral chrompodellid species likely possessed SMC5 and SMC6 and the loss of the two SMC subfamilies occurred at least on the branch leading to *Chromera velia*. We are currently unsure whether an undescribed chrompodellid “symbiont X” truly lacks SMC5/6, as the transcriptome was prepared from a limited number of the parasite/symbiont cells manually isolated from the host animal (Janouškovec et al. 2019). Thus, the library for sequencing may have missed the transcripts encoding SMC5 and SMC6.

Overall, the qualities/quantities of the ciliate transcriptome data may not be high enough to identify all of the six SMC subfamilies. We succeeded in identifying a set of SMC1-4 only in four out of the 15 species examined additionally (Fig. 2B). Nevertheless, we could find SMC5 or SMC6 in none of the 15 species examined additionally. Based on the surveys of SMC5/6 in the transcriptome data together with those in the three high-quality genome data, we propose that the common ancestor of ciliates lost both SMC5 and SMC6 secondarily.

### Phylogenetic relationship among the SMC subfamilies in eukaryotes: Rooted analyses

We subjected only SMC sequences that cover both highly conserved “block” at the N-terminus (i.e., Walker A and Q-loop motifs) to those at the C-terminus (i.e., Signature and Walker B motifs) to the ML and Bayesian phylogenetic analyses, along with the bacterial and archaeal SMC sequences, and Rad50/SbcC sequences (Rad50 in Bacteria is termed as SbcC). As the ML and Bayesian phylogenies agreed with each other overall, and thus Bayesian posterior probabilities (BPPs) for the nodes of our interest are shown on the ML tree (Fig. 3A; the Bayesian consensus tree with BPPs is provided as Fig. S3). In the ML analysis, the six SMC subfamilies in eukaryotes formed individual clades with ML nonparametric bootstrap support values (MLBPs) ranging from 95-100%, ultra-fast bootstrap support values (UFBPs) ranging from 95-100%, and BPPs of 1.0 (nodes 1, 2, 4, 5, 8, and 9 in Fig. 3A; see Fig. S3 for the Bayesian consensus tree with BPPs). The clade of bacterial SMC homologs and that of Rad50/SbcC homologs were supported by MLBPs/UFBPs/BPPs of 75%/100%/1.0 and 94%/100%/1.0, respectively. The monophyly of the archaeal SMC (aSMC) homologs was not recovered (see below). For the ML and Bayesian trees with the full taxon names, we supply the treefiles as a part of Supplementary materials.

**Figure 3.**
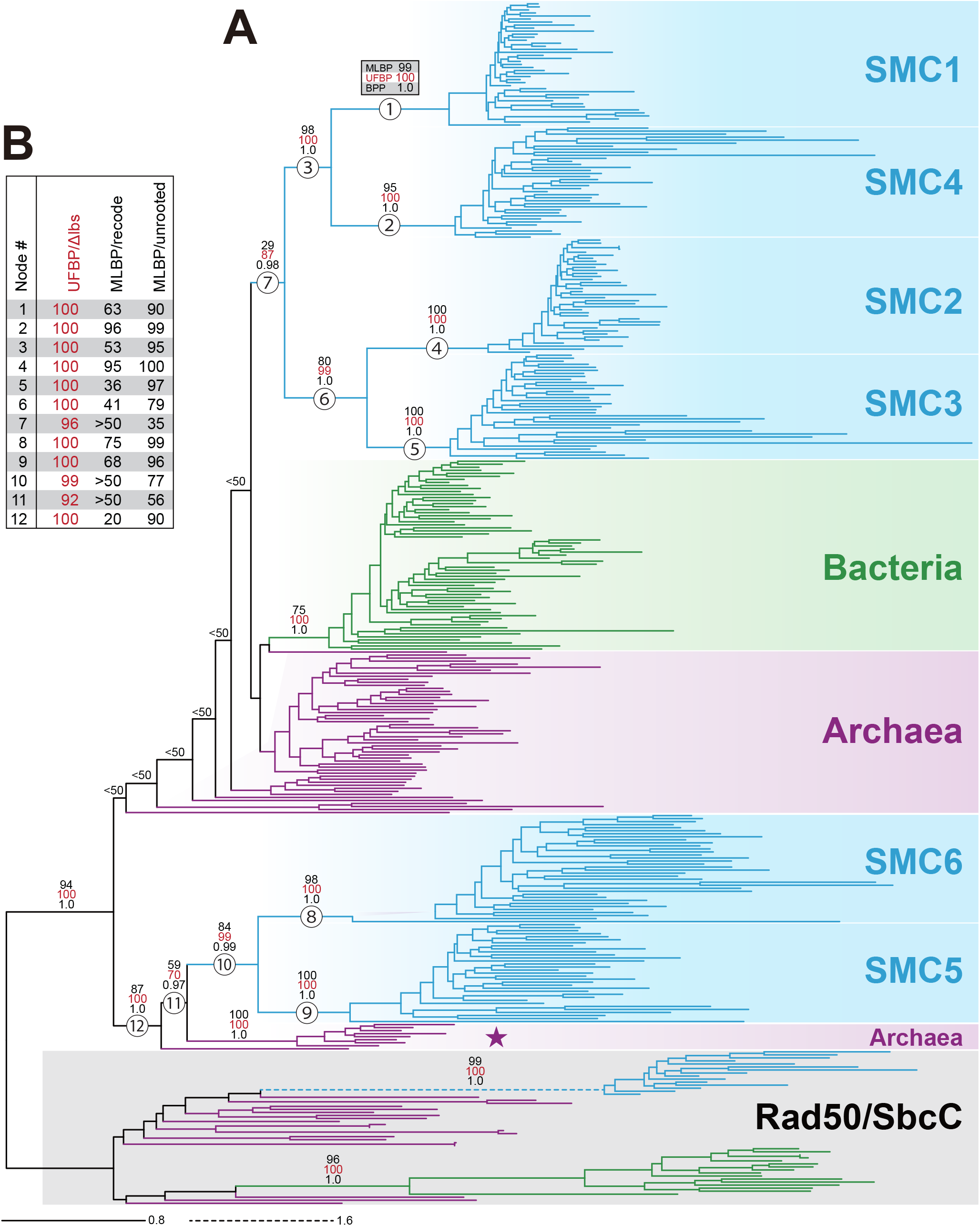
Maximum likelihood (ML) phylogeny of SMC sequences in Bacteria, Archaea, and Eukaryota rooted by Rad50/SbcC sequences. (A) Rooted ML phylogeny. All of the sequence names are omitted. The bacterial, archaeal, and eukaryotic branches are colored in green, purple, and blue, respectively. The clade of Rad50/SbcC sequences is shaded in grey. The ML nonparametric bootstrap support values (MLBPs), ultrafast bootstrap support values (UFBPs), and Bayesian posterior probabilities (BPPs) are displayed for the nodes that are critical to the evolution of the six SMC subfamilies of eukaryotes. Many of the deep splits received MLBPs smaller than 50% (labeled as “<50”). The 9 archaeal branches are highlighted by a star as “SMC5/6-related archaeal SMC homologs” (see the main text for the details). (B) Summary of the statistical support for nodes 1-12. The UFBPs in the first column and MLBPs in the second column were calculated from the SMC/SMC-related alignment without extremely long-branch sequences (UFBP/Δlbs) and that processed by the procedure recoding 20 amino acids into 4 bins (MLBP/recode), respectively. The MLBPs in the last column were calculated from the alignment from which Rad50/SbcC sequences were excluded (MLBP/unrooted).

In the SMC phylogeny presented in Cobbe and Heck (2004), SMC1 and SMC4, SMC2 and SMC3, and SMC5 and SMC6 formed the SMC1+4 clan, the SMC2+3 clan, and the SMC5+6 clan, respectively, although no ML bootstrap analysis was carried out. In contrast, we subjected the alignment containing SMC and Rad50/SbcC in Bacteria, Archaea, and Eukaryota to the ML, ML bootstrap, and Bayesian analyses, and the SMC1+4 clan, SMC2+3 clan, and SMC5+6 clan recovered with MLBPs/UFBPs/BPPs of 98%/100%/1.0, 80%/99%/1.0, and 84/99%/0.99, respectively (nodes 3, 6, and 10 in Fig. 3A). The SMC1+4 and SMC2+3 clans grouped together, albeit the nodes uniting SMC1-4 received an MLBP/UFBP/BPP of 29%/87%/0.98 (node 7 in Fig. 3A). Surprisingly, the SMC5+6 clan showed an intimate affinity to a subset of aSMC homologs (highlighted by a star in Fig. 3A), instead of other eukaryotic SMC subfamilies.

The clade of the aSMC homologs found in 8 metagenomes, which belong to Crenarchaota, Candidatus Verstraetearchaeota, Thaumarchaeota, and Candidatus Bathyarchaeota, joined the SMC5+6 clan with an MLBP/UFBP/BPP of 59%/70%/0.97 (node 11 in Fig. 3A). Then, the SMC homolog found in the metagenome of an archaeon belonging to Korarchaeota was connected to the clade of the SMC5+6 clan and the 8 aSMCl homologs with an MLBP/UFBP/BPP of 87%/100%/1.0 (node 12 in Fig. 3A). The metagenomes, which carry the 9 aSMC homologs branched at the base of the SMC5+6 clan, belong commonly to the TACK superphylum.

We further pursued the phylogenetic relationship among SMC1-6. As the branches leading to the eukaryotic SMC subfamilies were long, we cannot exclude the possibility of the ML tree shown in Fig. 3A being biased by long-branch attraction (LBA) artifacts (Bergsten 2005). To examine the possibility mentioned above, we eliminated extremely long-branch sequences from the SMC/SMC-related alignment and repeated the ML phylogenetic analysis. However, the tree topology before and that after eliminating extremely long-branch sequences were essentially unchanged (Fig. S4 and the UFBPs for nodes 1-12 are shown in the column labeled as “UFBP/Δlbs” in Fig. 3B; for the ML tree with full taxon names, see the treefile supplied as a part of Supplementary materials). Thus, the extremely long-branch sequences unlikely biased the phylogenetic relationship among the six SMC subfamilies in eukaryotes.

The SMC/SMC-related sequences are divergent (Fig. 3A), implying that the amino acid substitutions in many alignment positions have been saturated. We here recoded 20 amino acids in the SMC/SCM-related alignment into four bins to ameliorate substitution saturation which potentially causes various artifacts in tree reconstruction including LBA (Susko and Roger 2007). In the ML tree inferred from the recoded alignment, the monophyly of SMC1-6 was not recovered and, overall, the MLBPs for the nodes of interest were lower than the corresponding values calculated from the original alignment (Fig. S5 and the MLBPs for nodes 1-12 are shown in the column labeled as “MLBP/recode” in Fig. 3B; for the ML tree with full taxon names, see the treefile supplied as a part of Supplementary materials). The SMC1+4 and SMC2+3 clans were separated from each other by the SMC homologs of bacteria and a subset of archaea, albeit the nodes separating the SMC1+4 and SMC2+3 clans were not supported by MLBPs greater than 50% (Fig. S5). The ML phylogeny inferred from the original alignment (node 7 in Fig. 3A) and that from the recoded alignment (Fig. S5) are not contradictory to each other, as both left a large room for the relationship between the SMC1+4 and SMC2+3 clans. The statistical support for the SMC5+6 clan was also reduced by the recoding procedure. The 9 aSMC homologs, which were placed at the base of the SMC5+6 clan in the first ML analysis (marked by a star in Fig. 3A), formed a clade and branched directly with the SMC6 clade (Fig. S5). The SMC5 clade was then connected with the clade of SMC6 plus the 9 aSMC homologs. Nevertheless, the above-mentioned nodes were poorly supported (MLBPs <50%; Fig. S5; see also node 12 in Fig. 3B). In sum, we conclude that the monophyly of SMC1-6 was not positively supported in the ML analysis of the recoded alignment.

Altogether, it is unlikely that the phylogenetic relationship among the six SMC subfamilies in eukaryotes was biased significantly by the LBA artifact which stemmed from long-branch sequences or substitution saturation in the original alignment.

### Phylogenetic relationship among the SMC subfamilies in eukaryotes: Unrooted analyses

We then conducted an extra set of ML analyses excluding the Rad50/SbcC (outgroup) sequences. As the SMC and Rad50/SbcC homologs are distantly related to each other (Fig. 3A), the outgroup potentially introduced LBA artifacts, which hindered the recovery of the monophyly of the six SMC subfamilies in eukaryotes, in the rooted analyses (e.g., Fig. 3A). Significantly, the phylogenetic relationship among the SMC subfamilies was almost identical before and after the exclusion of the outgroup (Figs. 4A, S6, and the MLBPs for nodes 1-12 are shown in the column labeled as “MLBP/unrooted” in Fig. 3B; for the unrooted ML tree with full taxon names, see the treefile supplied as a part of Supplementary materials). Thus, the inclusion of the Rad50/SbcC homologs in the original alignment most likely gave no substantial impact on the SMC phylogeny.

**Figure 4.**
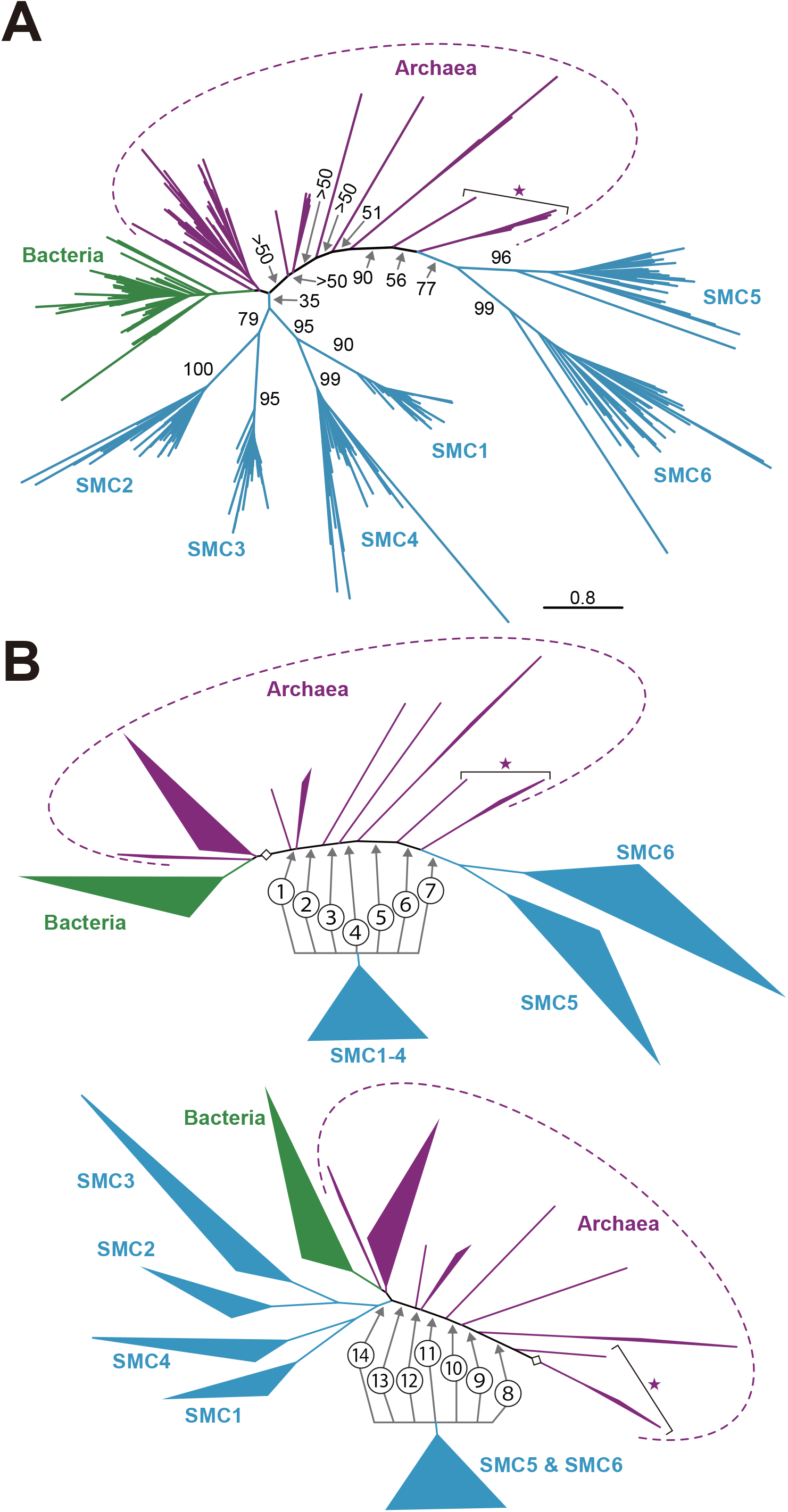
(A) Unrooted maximum likelihood (ML) phylogeny of SMC sequences in Bacteria, Archaea, and Eukaryota. All of the sequence names are omitted. The color scheme for bacterial, archaeal, and eukaryotic SMC branches is the same as described in the legend of Fig. 4. Only ML nonparametric bootstrap values for the major nodes are shown. (B) The scheme how to prepare the alternative trees for an approximately unbiased test. From the unrooted ML tree (see A), the first set of 7 alternative trees (Trees 1-7) were generated by pruning and regrafting of the clade of SMC1-4 into alternative positions labeled as 1-7 (see the upper figure). The second set of 7 alternative trees (Trees 8-14) were generated by pruning and regrafting of the clade of SMC5 and SMC6 into alternative positions labeled as 8-14 (see the lower figure). In both upper and lower figures, the diamonds represent the original position of the clade of SMC1-4 and that of SMC5 and SMC6 in the ML tree (Tree 0). In all figures, “SMC5/6-related archaeal SMC homologs” are highlighted by a star (see the main text for the details).

Neither rooted nor unrooted ML phylogeny favored the monophyly of the six SMC subfamilies in eukaryotes (Figs. 3A and 4A). We here examine the alternative trees bearing the monophyly of SMC1-6 by an approximately unbiased (AU) test (Shimodaira 2002). To avoid any potential biases in tree reconstruction caused by the outgroup, we conducted the AU test based on the unrooted alignment. If we discard the hypothesis assuming the monophyly of SMC1-6 in the test based on the unrooted alignment, SMC1-6 cannot form the clade in the rooted phylogeny. For the AU test, we modified the unrooted ML tree (Fig. 4A) by pruning and regrafting the subtree containing both SMC1+4 and SMC2+3 clans in 7 alternative positions, as illustrated in the upper part in Fig. 4B. Likewise, the SMC5+6 clan was pruned from the original position and regrafted to 7 alternative positions to generate an extra 7 alternative trees (the lower part in Fig. 4B). Only two out of the 14 alternative trees (i.e., Trees 3 and 8) failed to be rejected at a 5% α-level (Table 1). Significantly, Trees 7 and 14, which support the monophyly of SMC1-6, were rejected at a 1% α-level (*P* values of 3.49 × 10^−35^ and 0.00961, respectively; Table 1).

**Table 1.**
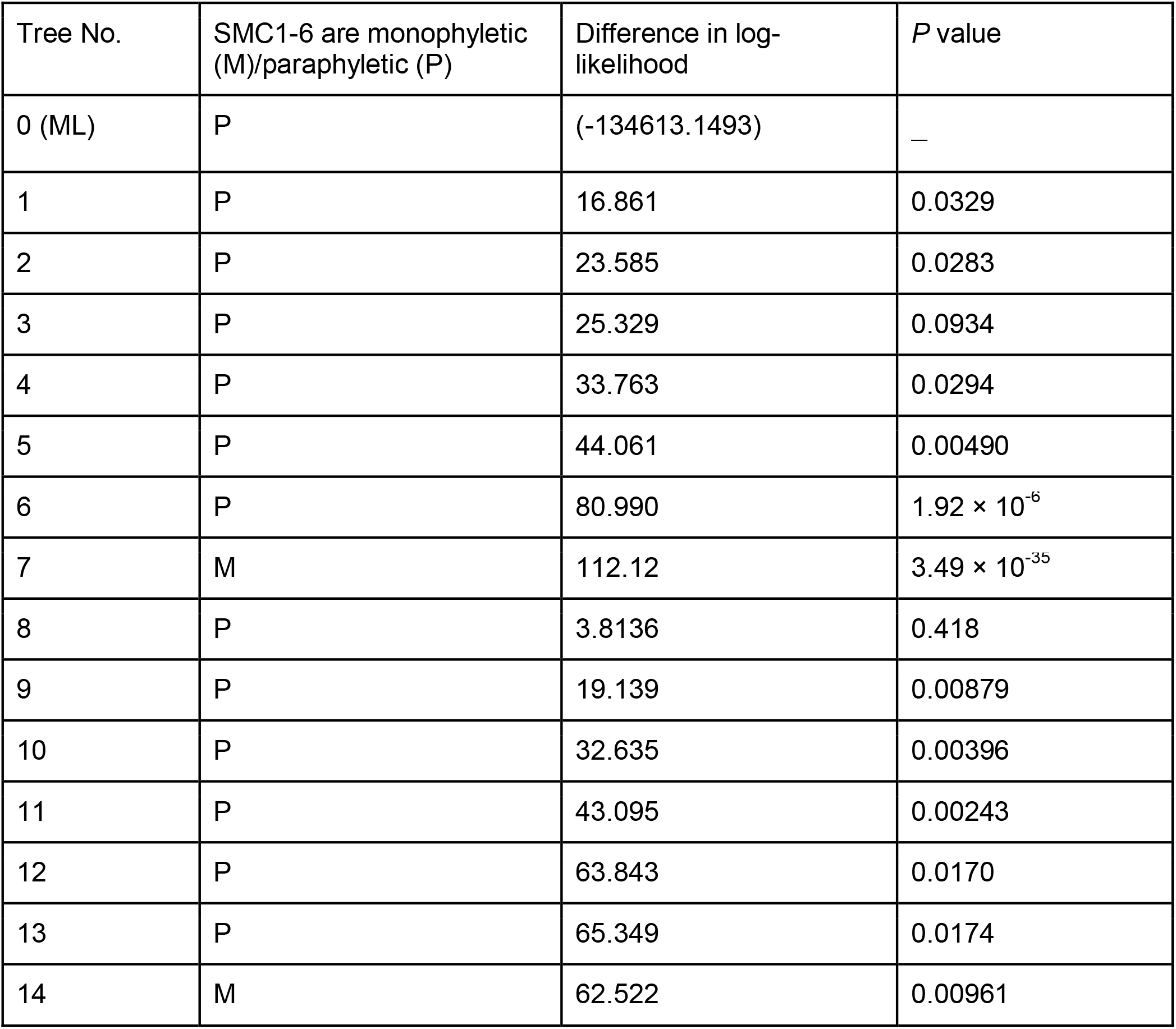
An AU test assessing the phylogenetic relationship among SMC1-6.

| Tree No. | SMC1-6 are monophyletic (M)/paraphyletic (P) | Difference in log-likelihood | *P* value |
| --- | --- | --- | --- |
| 0 (ML) | P | (-134613.1493) | — |
| 1 | P | 16.861 | 0.0329 |
| 2 | P | 23.585 | 0.0283 |
| 3 | P | 25.329 | 0.0934 |
| 4 | P | 33.763 | 0.0294 |
| 5 | P | 44.061 | 0.00490 |
| 6 | P | 80.990 | $1.92 \times 10^{-6}$ |
| 7 | M | 112.12 | $3.49 \times 10^{-35}$ |
| 8 | P | 3.8136 | 0.418 |
| 9 | P | 19.139 | 0.00879 |
| 10 | P | 32.635 | 0.00396 |
| 11 | P | 43.095 | 0.00243 |
| 12 | P | 63.843 | 0.0170 |
| 13 | P | 65.349 | 0.0174 |
| 14 | M | 62.522 | 0.00961 |

Both standard phylogenetic analyses and the AU test consistently suggested that the SMC5+6 clan is evolutionarily distantly related from the rest of the SMC subfamilies in eukaryotes (i.e., SMC1-4).

### Phylogenetic diversity of the archaeal SMC

The rooted and unrooted SMC phylogenies indicated that the 9 aSMC homologs, all of which were found in the metagenomes belonging to the TACK superphylum, grouped with SMC5 and SMC6 with high statistical support (highlighted by stars in Figs 3A and 4A; henceforth here, designated as “SMC5/6-related aSMC homologs”). Among the 512 aSMC homologs retrieved initially from the GenBank nr database (see Materials & Methods), we found 80 SMC5/6-related aSMC homologs including the 9 homologs described above (Fig. S7; for the ML tree with full taxon names, see the treefile supplied as a part of Supplementary materials). We here propose to split the 512 aSMC homologs into two categories. The first aSMC category contains the 432 homologs that show no phylogenetic affinity to SMC5 and SMC6, and was found in phylogenetically broad species/metagenomes belonging to TACK superphylum, Euryarchaeota, DPANN group, Asgard group, and Candidatus Thermoplasmatota (designated as the “canonical aSMC”; Fig. 5A). Considering the phylogenetic distribution, the canonical aSMC likely has been inherited from the ancestral archaeon to the major descendents. The second category comprises SMC5/6-related aSMC homologs found in the restricted set of the species/metagenomes. 66 out of the 80 SMC5/6-related aSMC homologs identified here were found in the species/metagenomes of the TACK superphylum (Fig. 5B). However, the species/metagenomes bearing SMC5/6-related aSMC homologs appeared to span Euryarchaeota, Asgard group, and Candidatus Thermoplasmatota, implying the early emergence of SMC5/6-related aSMC prior to the divergence of major archaeal lineages. Although no SMC5/6-related aSMC homolog was found in any member of the DPANN group, we anticipate finding some members carrying SMC5/6-related aSMC in this lineage in the future. The distributions of the canonical and SMC5/6-related aSMC in Archaea force us to assume the ancestral archaeon with the two distinct SMC homologs. Intriguingly, 44 out of the 80 species/metagenomes were found to possess both canonical and SMC5/6-related aSMC homologs and may represent the SMC repertory in the ancestral archaeon. As the majority of the species/metagnomes investigated here possesses either canonical or SMC5/6-related aSMC homolog, the loss of one of the two distinct SMC homologs most likely occurred at a high rate in the tree of Archaea.

**Figure 5.**
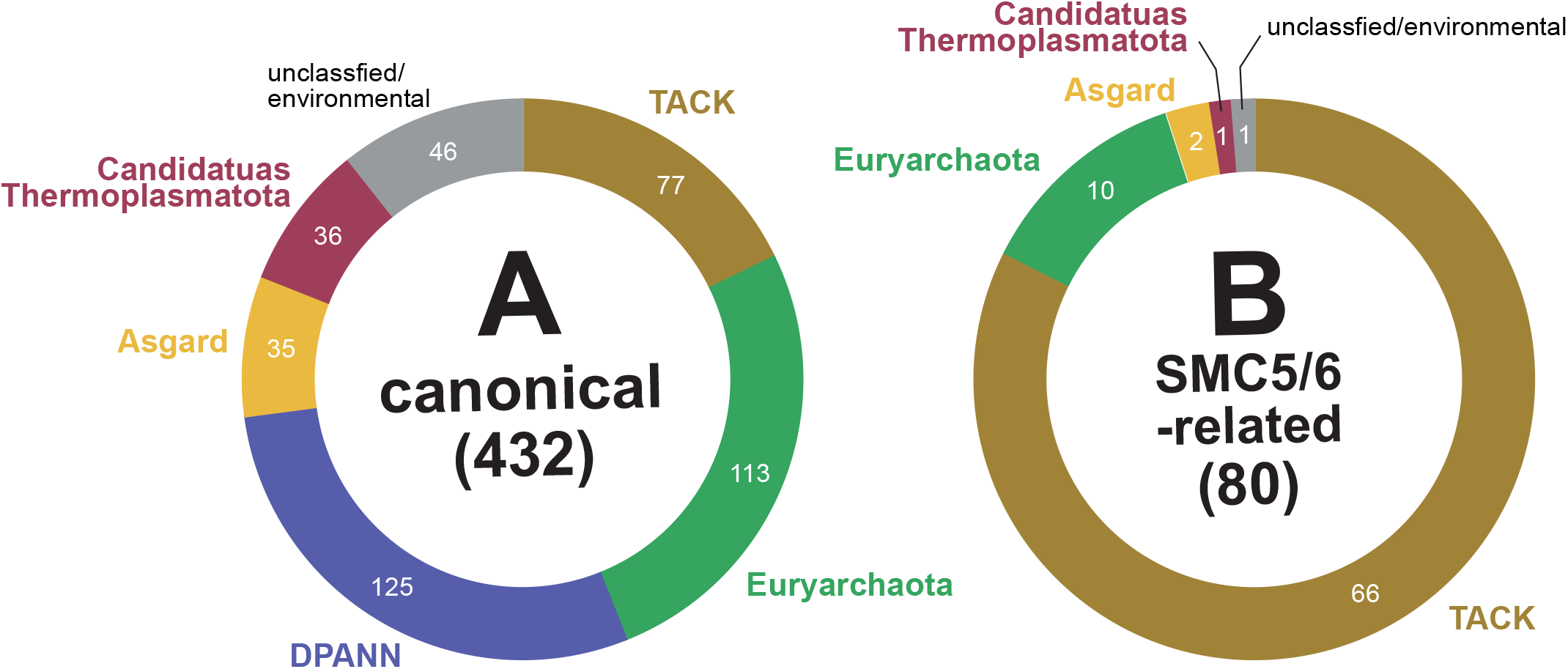
Taxonomic classification of the archaeal species/metagenomes encoding 512 SMC homologs retrieved from the GenBank database. (A) Archaeal SMC homologs that showed no phylogenetic affinity to SMC5 and SMC6 (“canonical aSMC”; see the main text). (B) Archaeal SMC homologs related to SMC5 and SMC6 (“SMC5/6-related aSMC”; see the main text). The taxonomic classification was followed by the NCBI Taxonomy database.

## DISCUSSION

### LECA possessed a set of cohesin, condensin, and the SMC5/6 complex

The pioneering work by Cobbe and Heck (2004) proposed the ubiquity of SMC1-6 among eukaryotes, albeit their conclusion was drawn from the survey against a phylogenetically restricted set of eukaryotes (i.e., 16 opisthokonts, two land plants, and two trypanosomatids). Transcriptome and/or genome data are now available for phylogenetically diverse eukaryotes so that the current study could search SMC homologs in a much broader diversity of eukaryotes than Cobbe and Heck (2004). We subjected 59 species, which represent all of the currently recognized major branches in the tree of eukaryotes plus “orphan” species. Fortunately, the search conducted in this study confirmed the ubiquity of SMC1-6 among eukaryotes (Fig. 1). We firmly conclude that the six SMC subfamilies had already existed in the genome of the LECA and the LECA most likely used cohesin, condensin, and the SMC5/6 complex for the maintenance of chromosomes.

Besides SMC1-6, non-SMC proteins (accessory subunits) comprise cohesin, condensin, and the SMC5/6 complex but their phylogenetic distributions have not been examined prior to this study. We searched for Rad21/Scc1 and STAG1/Scc3 in cohesin, CP-D2, CAP-G, and CAP-H in condensin, and Nse-1-4 in the SMC5/6 complex, in the 59 species in this study (Fig. S1). Due to the relatively low conservation at the amino acid sequence level, the search of the accessory subunits was less successful than that of the SMC proteins with highly conserved sequence motifs for ATP-binding. Nevertheless, we found the accessory subunits across the tree of eukaryotes. The phylogenetic analysis of the kleisin superfamily, namely Rad21/Scc1, CAP-H, Nse4, and ScpA in bacteria and archaea, suggested that the eukaryotic kleisins share a common ancestry (Fig. S2). The results described above suggest that the subunit compositions of cohesin, condensin, and the SMC5/6 complex of the LECA were similar to those of modern eukaryotes.

Although the early emergence of cohesin, condensin, and the SMC5/6 complex in the eukaryotic evolution (see above), we unveiled the secondary losses of SMC5 and SMC6 on separate branches in the tree of eukaryotes. By combining the survey of SMC1-6 in 59 phylogenetically diverse eukaryotes plus additional 42 species, we propose that secondary loss of SMC5 and SMC6 occurred in (i) the common ancestor of kinetoplastids and diplonemids, (ii) that of dinoflagellates and perkinsids, (iii) that of ciliates, and (iv) on the branch leading to the chrompodellid *Chromera velia* (Figs. 2A &2B). Importantly, our results imply that the SMC5/6 complex is not absolutely indispensable for cell viability. If so, extra eukaryotic groups/species lacking SMC5 and SMC6 likely have been overlooked among those of which large-scale sequence data are currently not available, and novel ones that are currently unknown to science.

### Multiple origins of eukaryotic SMC subfamilies

The majority of bacteria and archaea possess a single type of SMC, suggesting that the two identical SMC proteins form a homodimeric complex (Soppa 2001). In contrast, eukaryotes possess the six SMC subfamilies to constitute the three distinct heterodimeric complexes, with exception of some lineages lacking SMC5 and SMC6 (see above). It has been assumed that the SMC subfamilies in eukaryotes emerged from a single ancestral SMC through multiple rounds of gene duplication followed by functional divergence (Cobbe and Heck 2000; Cobbe and Heck 2004). To retrace how the repertory of the SMC subfamilies has been shaped prior to the emergence of the LECA, the phylogenetic relationship of the SMC subfamilies in Bacteria, Archaea, and Eukaryota is critical.

To phylogenetically examine the evolution of SMC in the three domains of life, SMC-related proteins (e.g., Rad50/SbcC), which had been present prior to the three domains, are indispensable as the outgroup. If all of the six SMC subfamilies in eukaryotes emerged from a single ancestral SMC protein, the monophyly of SMC1-6 should be recovered by excluding the bacterial and archaeal homologs in the rooted SMC phylogeny. Curiously, as shown in Fig. 3A, the clade of SMC1-4 and the SMC5+6 clan were separated from each other by the bacterial and archaeal SMC homologs plus the outgroup sequences. It is noteworthy that particular archaeal homologs showed an intimate phylogenetic affinity to SMC5 and SMC6 (highlighted by a star in Fig. 3A; see below). The exclusion of extremely long branches, that of the outgroup sequences, or recoding of the 20 amino acids into four bins introduced essentially no change in the distant relationship between SMC1-4 and SMC5 and SMC6, suggesting that LBA artifacts, if exist, were negligible in the SMC phylogeny (Figs. 3A, 3B, 4A, and S4-6). Moreover, an AU test rejected the alternative trees bearing the monophyly of SMC1-6 at a 1% α-level (Trees 7 and 14; Fig. 4B and Table 1). Thus, we conclude that the clade of SMC1-4 and that of SMC5 and SMC6 unlikely grouped together directly, regardless of the presence/absence of the outgroup. Overall, the results described above consistently and strongly suggest the aSMC homolog most closely related to SMC1-4 is distinctive from that most closely related to SMC5 and SMC6. Here we propose that SMC1-4 and SMC5 and SMC6 were derived separately from the canonical aSMC and SMC5/6-related aSMC, respectively.

### SMC evolution in Archaea and Eukaryota

The SMC phylogeny suggested that SMC1-4 and SMC5 and SMC6 emerged from distinct aSMC homologs (see above), suggesting that the diversity and evolution of SMC in Archaea and those in Eukaryota are intimately related to each other. Our survey of the SMC homologs in Archaea suggested that the two distinct aSMC homologs, namely canonical and SMC5/6-related aSMC homologs, existed prior to the divergence of major lineages in Archaea (Fig. 6A). We here propose a massive number of secondary losses of one of the two aSMC homologs in the evolution of Archaea, resulting in only a minor fraction of the modern archaea keeping both homologs. In the archaea bearing both canonical and SMC5/6-related aSMC homologs, the two proteins likely constituted two distinct homodimeric SMC complexes, rather than a single heterodimeric complex. The loss of one of the two SMC proteins likely occurred frequently in the tree of Archaea but, if the redundancy in SMC complexes existed, should not have resulted in severe damage to the cell viability.

**Figure 6.**
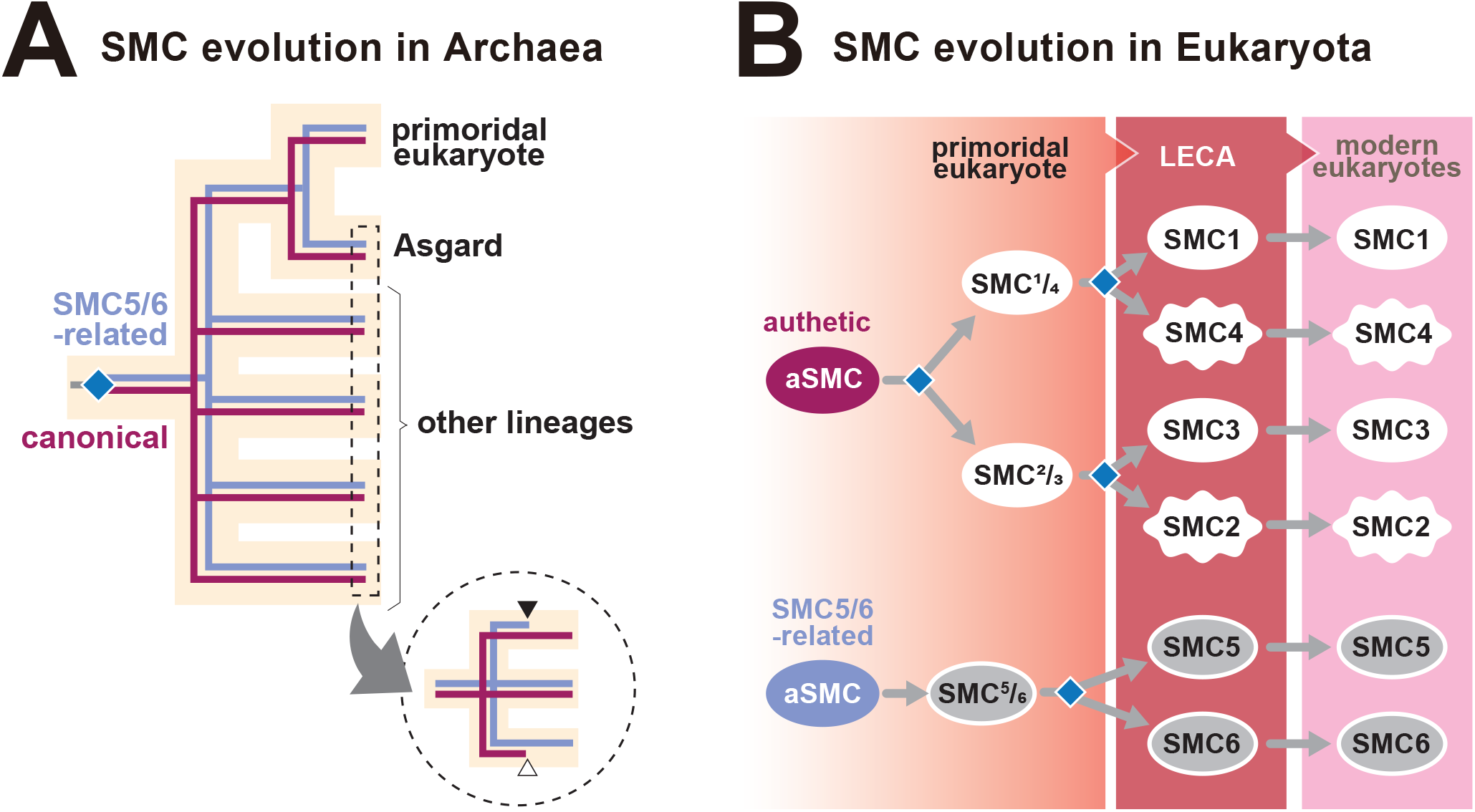
A scenario for the evolution of SMC in Archaea and Eukaryota. (A) SMC evolution in Archaea. We assume that the ancestral archaeon possessed two distinct types of SMC, one is shared the majority of modern archaea and ancestral to SMC1-4 in eukaryotes (labeled as “canonical”; colored in dark red) and the other is found in a restricted set of modern archaea and closely related to SMC5 and SMC6 in eukaryotes (labeled as “SMC5/6-related”; colored in pale blue). The two types of SMC emerged by the duplication of a SMC gene in the ancestral archaeon (shown by a diamond) and were inherited to major archaeal lineages and the primordial eukaryote which is closely related to the Asgard group. As demonstrated in the circle, parallel secondary losses of one of the two aSMC homologs occurred during the divergence of each archaeal lineage. Closed and open arrowheads represent the loss of SMC5/6-related aSMC and that of the canonical aSMC, respectively. (B) SMC evolution in Eukaryota. We hypothesize that SMC1-6 in modern eukaryotes (the right column) were descended from the last eukaryotic common ancestor (LECA) (the center column). Because of the phylogenetic affinity between SMC1 and SMC4 and that between SMC2 and SMC3 (see Fig. 3A), the ancestral molecule for SMC1 and SMC4 and that for SMC2 and SMC3 may have existed in the primordial eukaryote (“SMC¼” and “SMC⅔”; the left column). Prior to the LECA, two separate gene duplication events followed by functional divergence (shown by diamonds) yielded SMC1-4. We assume that one of the two aSMC homologs (i.e., “canonical” SMC) gave rise to SMC¼ and SMC⅔ via gene duplication followed by functional divergence (shown by a diamond). In addition to SMC¼ and SMC⅔, the primordial eukaryote likely possessed “SMC□,” which was derived from an SMC5/6-related aSMC in Archaea. During the transition from the primordial eukaryote to the LECA, SMC□ diverged into SMC5 and SMC6 via gene duplication followed by functional diversification (shown by a diamond).

The first eukaryote likely emerged from one of the archaeal lineages, which retains both canonical and SMC5/6-related aSMC homologs. Unfortunately, the SMC sequence data had no sufficient phylogenetic resolution to pinpoint the precise species/lineage whose aSMC homologs gave rise to SMC1-4 and SMC5 and SMC6 in the LECA. The difficulty in resolving deep splits in the SMC phylogeny likely stems from the large difference in tempo and mode of amino acid substitution among the SMC sequences in Eukaryota and Archaea.

Nevertheless, considering our current belief of the Asgard origin of Eukaryota (Zaremba-Niedzwiedzka et al. 2017; Williams et al. 2020), it is reasonable to nominate the canonical and SMC5/6-related aSMC proteins in a member of the Asgard group as the ancestral molecule of SMC1-4 and that of SMC5 and SMC6 in the LECA (and its descendants). To bridge the gap in the repertory of SMC between Archaea and Eukaryota, we here introduce that the “primordial eukaryote,” which existed prior to the LECA and used at least three distinct SMC proteins, (i) the ancestral protein for SMC1 and SMC4, (ii) that for SMC2 and SMC3, and (iii) that for SMC5 and SMC6 (henceforth designated as “SMC¼,” “SMC⅔,” and “SMC□,” respectively; Fig. 6B). SMC¼ and SMC⅔ were yielded by a single duplication of the canonical aSMC gene followed by functional divergence (a diamond in Fig. 6B), while SMC□ was the direct descendant of SMC5/6-related aSMC homologs. The primordial eukaryote likely possessed two SMC complexes, one is a heterodimeric complex constituted by SMC¼ and SMC⅔ and the other is a homodimeric complex of two SMC□ proteins. Then, two separate but concomitant gene duplications followed by functional divergence generated SMC1-4 (diamonds in Fig. 6B) yielding two heterodimeric SMC complexes (i.e., cohesin and condensin) in the LECA. The duplication of SMC□ gene followed by function divergence was also required to yield the third heterodimeric SMC complex (i.e., SMC5/6 complex) en route from the primordial eukaryote to the LECA.

## CONCLUSIONS

In this study, we reexamined the ubiquity and evolution of the six eukaryotic SMC subfamilies based on the transcriptome and genome data accumulated from phylogenetically diverse eukaryotes. The survey of SMC sequences conducted in this study confirmed the ubiquity of SMC1-6 in eukaryotes (Fig. 1). Significantly, we here revealed two novel aspects of the SMC evolution in eukaryotes. First, we identified multiple secondary losses of SMC5 and SMC6 in the tree of eukaryotes (Figs. 2A & 2B), questioning their absolute necessity for cell viability. Second, we noticed that SMC5 and SMC6, and SMC1-4 were most likely derived from distinct ancestral molecules in Archaea (Figs. 3A, 3B, 4A, and Table 1). We finally proposed that the first eukaryote inherited two distinct SMC homologs from an archaeon (likely a member of the Asgard group) and then two separate but concomitant duplications of one of the two SMC genes to yield SMC1-4, while SMC5 and SMC6 emerged through a single duplication of the other SMC gene (Figs. 6A and 6B).

## MATERIALS AND METHODS

### Surveys of SMC proteins in Eukaryota

To examine the ubiquity of SMC1-6 in eukaryotes, we searched for SMC amino acid sequences in 59 phylogenetically diverse species. For 36 out of the 59 eukaryotes, the SMC sequences were searched for in NCBI GenBank non-redundant (nr) protein sequences database by BLASTP (Camacho et al. 2009). For the rest of the 23 eukaryotes, the SMC sequences were retrieved from the corresponding transcriptome data. The searches against the GenBank nr database and those against the contig data assembled from the transcriptome data are separately explained below.

For each of the 36 species, a BLASTP search was conducted by using the amino acid sequences of human SMC homologs as queries (GenBank accession numbers are Q14683 for SMC1, O95347 for SMC2, Q9UQE7 for SMC3, Q9NTJ3 for SMC4, Q8IY18 for SMC5, and Q96SB8 for SMC6). The amino acid sequences, which showed high similarity to the human SMC1-6 in the initial BLAST analyses, were subjected to reciprocal BLASTP against the GenBank nr protein database. We retained the sequences, which showed similarities to the SMC sequences deposited in the database in the second BLAST analyses, as the candidates for the SMC sequences.

For the 23 eukaryotes, we downloaded their transcriptome data from NCBI Sequence Read Archive (Sayers et al. 2020) to identify the transcripts encoding SMC proteins. The raw sequence reads were trimmed by fastp v0.20.0 (Chen et al. 2018) with the -q 20 -u 80 option and then assembled by Trinity v2.8.5 or v2.10.0 (Grabherr et al. 2011). The candidate sequences of SMC proteins were retrieved from the assembled data by TBLASTN with human SMC1-6 as queries followed by reciprocal BLASTP against the GenBank nr protein database.

If a subset of SMC1-6 were not detected in the surveys described above, we switched the queries from the human homologs to those sampled from the close relatives of the target organisms and repeated the BLAST survey.

The search of SMC proteins in the 59 eukaryotes revealed the secondary loss of SMC5 and SMC6 in phylogenetically distinct taxonomic groups, namely kinetoplastids, dinoflagellates plus *Perkinsus marina*, ciliates, and *Chromera velia* (see the Results). Thus, we additionally assessed the repertories of SMC1-6 in the phylogenetic relatives of the above-mentioned species (42 species in total).

For most of the 42 species, only transcriptomic data were available. Thus, the survey of SMC sequences against the transcriptome data was repeated as described above. For the additional survey of SMC1-6 in Euglenozoa including kinetoplastids, diplonemids, and euglenids, we used two sets of queries, namely human SMC1-6 and *Naegleria fowleri* SMC1-6. Likewise, *Vitrella brassicaformis* SMC1-6, together with the human homologs, were used as queries for the additional survey in Alveolata including apicomplexans, chromopodellids, dinoflagellates, and ciliates.

### Surveys of SMC proteins in Archaea and Bacteria

The amino acid sequences of bacterial and archaeal SMC homologs were retrieved from the GenBank refseq and nr protein databases, respectively (see below). First, we searched for the archaeal proteins belonging to the SMC superfamily by using BLASTP (taxid = 2157; max_target_seq = 50,000). The conserved N- and C-terminal portions of the following sequences were subjected separately to BLASTP analyses as the queries; human SMC1-6, a single bacterial SMC (WP_045231317.1), three aSMCs (TFG22733.1; AAB89690.1; WP_148681221.1), a single MukB (NWA44283.1), a single SbcC (OYY36098.1), and a single RecN (WP_187956897.1). We pooled the GenBank entries matched to the individual queries mentioned above with *E*-values smaller than 10^−5^ in the initial set of BLASTP analyses and further selected the entries, of which sequence lengths ranged between 500 and1500 amino acid residues. After dissolving the redundancy in the GenBank entries retrieved by the BLASTP analyses with the multiple queries (see above), we obtained 2,363 entries that cover the entire proteins belonging to the SMC superfamily. Then, the amino acid sequences of the selected entries were subjected to cd-hit (-c 0.6) to reduce the redundancy at the amino acid sequence level, leaving 559 sequences for a preliminary phylogenetic analysis. The 559 aSMC candidate sequences were aligned with the previously identified SMC sequences in Eukaryota, Archaea, and Bacteria, as well as RecN, MukB, and Rad50/SbcC sequences (for the details of the alignment preparation, see below). We removed the alignment positions corresponding to the coiled-coil portion manually and trimmed the positions including gaps with trimal -gt 0.8. The alignment of 621 sequences including the 559 aSMC candidates was phylogenetically analyzed with the ML method with the LG + Γ + I + F model by IQ-TREE v1.6.12 (Nguyen et al. 2015). Based on the preliminary phylogenetic analysis, 36 out of the 559 aSMC candidates appeared to be Rad50, MukB, or RecN. The phylogenetic analysis also grouped 11 aSMC candidates tightly with the bacterial SMC homologs. Thus, we regarded the remaining 512 sequences as the genuine aSMC homologs. We finally selected 63 sequences, which represent the SMC diversity in Archaea, from the 512 sequences identified by the above-mentioned procedure, together with the previously identified sequences. The accession numbers of the 512 aSMC amino acid sequences are summarized in a spreadsheet file supplied as a part of Supplementary materials.

To retrieve diverse bacterial SMC (bSMC) sequences, we repeated the BLASP analysis described above against the refseq protein database (taxid = 2; max_target_seq = 50,000) with the same set of queries described above except only *Archaeoglobus fulgidus* sequence (AAB89690.1) was considered as the archaeal aSMC query). 11,534 GenBank entries, which cover the entire proteins, were retrieved. We further selected 1,742 sequences by reducing the redundancy at the amino acid sequence level. The preliminary alignment of the 1,742 bSMC candidates and previously identified SMC, RecN, MukB, and Rad50/SbcC sequences (1,871 in total) were prepared and phylogenetically analyzed as described above. By assessing the preliminary ML analysis, we excluded 1,299 non-SMC sequences, leaving 443 bSMC homologs. Finally, 63 sequences, which represent the bSMC diversity, were selected from the 443 bSMC sequences for the phylogenetic analyses (see below).

### Surveys of Rad50/SbcC proteins

Several proteins related to SMC (SMC-related proteins) have been known in Bacteria, Archaea, and/or Eukaryota. For instance, Rad50 is one of the SMC-related proteins conserved among the three domains of life. We retrieved Rad50 amino acid sequences of 15 eukaryotes and 20 archaea from the NCBI GenBank nr database by the BLAST search as described above. The BLAST searches for eukaryotic and archaeal Rad50 were separately conducted with the human and *Archaeoglobus fulgidus* homologs (GenBank accession numbers Q92878 and NC_000917.1) as queries, respectively. The amino acid sequences of SbcC of 16 bacteria were retrieved from the GenBank database through keyword searches.

### Phylogenetic analyses

We newly identified the candidate sequences for SMC proteins sampled from phylogenetically broad eukaryotes (see above). The SMC candidate sequences identified in this study and together with the previously known SMC sequences (typically those of animals, fungi, and land plants), were subjected to the ML phylogenetic analysis with the LG + Γ + I model. This preliminary analysis aimed to filter extremely divergent sequences and the sequences that could not be classified into any of the eukaryotic SMC sub-families with confidence. We examined the ubiquity of each of the six SMC subfamilies in eukaryotes based on the filtered SMC sequences (298 sequences in total).

To retrace the evolution of the six SMC subfamilies in eukaryotes, we further selected 240 SMC sequences containing both of the two ATP-binding motifs in the N- and C-termini from the 298 eukaryotic SMC sequences. After the exclusion of redundant sequences by CD-HIT (Fu et al. 2012), we retained 220 eukaryotic SMC sequences for the phylogenetic analyses described below.

Individual sub-families of eukaryotic SMC were separately aligned by MAFFT v7.453 (Katoh and Standley 2013) with the L-INS-I option. Likewise, 63 bSMC, 63 aSMC, and 51 Rad50/SbcC amino acid sequences were separately aligned as described above. The alignments of SMC1-6, bSMC, aSMC, and Rad50/SbcC sequences were combined into the “SMC/SMC-related” alignment by MAFFT with --merge option. The resultant alignment contained 35 of SMC1, 41 of SMC2, 37 of SMC3, 37 of SMC4, 34 of SMC5, and 36 of SMC6 sequences. Manual exclusion of ambiguously aligned positions, followed by trimming of gap-containing positions by trimAI v1.2 (Capella-Gutiérrez et al. 2009) with -gt 0.8 option, left 293 unambiguously aligned amino acid positions in 346 SMC and 51 Rad50/SbcC for the phylogenetic analyses described below. The accession numbers of the amino acid sequences included in the SMC/SMC-related alignment are summarized in a spreadsheet file supplied as a part of Supplementary materials.

We subjected the SMC/SMC-related alignment to the ML analysis with the LG + R9 + F + C60 model to infer the ML tree and calculate UFBPs. We also conducted a 100-replicate non-parametric ML bootstrap analysis with the LG + R9 + F + C60 + PMSF model (the guide tree was reconstructed with the ML method with the LG + R9 + F + C60 model). The ML tree search and non-parametric ML bootstrap analysis described above were repeated twice, once after excluding 9 extremely long-branch sequences and the other after excluding all of the Rad50/SbcC sequences. The original alignment was further modified by separating 20 amino acid characters into four bins containing (i) A, G, N, P, S, and T (amino acid characters are indicated by the one-letter code), (ii) C, H, W, and Y, (iii) D, E, K, Q, and R, and (iv) F, I, L, M, and V to ameliorate substitution saturation (Susko and Roger 2007). The recoded alignment was subjected to the ML tree search and 100-replicate ML non-parametric bootstrap analysis with the GTR + Γ + F model. IQ-TREE v1.6.12v2.1.2 (Nguyen et al. 2015) was used for all of the ML phylogenetic analyses described above.

The SMC/SMC-related alignment was also analyzed with Bayesian method by PhyloBayes v4.1 (Lartillot et al. 2009) using the CAT + GTR model. We ran two Markov chain Monte Carlo (MCMC) chains for 55,963 cycles and the first 14,000 cycles were discarded as burn-in. Although the indicator of the convergence among the MCMC chains remained large (maxdiff = 0.487909), we found that the two tree topologies of individual MCMC chains agreed to each other largely. Thus, the consensus tree with branch lengths and BPPs were calculated from the remaining trees.

We examined whether the six SMC subfamilies in eukaryotes are monophyletic by an AU test (Shimodaira 2002) considering the unrooted ML tree (Tree 0) and 14 alternative trees (Tree 1-14) as described below. In the unrooted ML tree (Fig. 4A), the clade of SMC1-4 and that of SMC5 and SMC5 were separated by the SMC homologs of bacteria and archaea. First, the clade of SMC1-4 was pruned from the original position in the ML tree and regrafted to the 7 nodes, which separated the two clades of the eukaryotic SMC subfamilies, to generate 7 alternative trees (Trees 1-7; the upper figure in Fig. 4B). Second, the pruning and regrafting of the clade of SMC5 and SMC6 was repeated to generate 7 alternative trees (Trees 8-14; the lower figure in Fig. 4B). We calculated site-wise log-likelihoods (site-lnLs) over each of Trees 0-14 and the resultant site-lnL data were subjected to an AU test. We applied the same substitution model applied to the unrooted ML phylogenetic analyses (see above). The site-lnL calculation and AU test were implemented in IQ-TREE v2.1.2 (Nguyen et al. 2015).

The 512 aSMC sequences retrieved from the GenBank (see “Surveys of SMC proteins in Archaea and Bacteria”) were aligned with 20 archaeal Rad50 sequences and 263 unambiguously aligned amino acid positions were subjected to the phylogenetic analysis described below. The ML tree with UFBPs was reconstructed from this “archaeal SMC/Rad50 alignment” with the LG + Γ + F + C20 model. The details of the preparation of the alignment and ML phylogenetic analysis are the same as described above.

### Analyses of accessory subunits of the SMC complexes in eukaryotes

Cohesin contains two accessory subunits Rad21/SCC1 and STAG1/SCC3, together with SMC1 and 3, while SMC2 and 4 interact with CAP-H, CP-D2, and CAP-G to constitute condensin. Nse1, Nse2, Nse3, Nse4, Nse5, and Nse6 are known to participate in the SMC5/6 complex. We searched for the accessory subunits in cohesin, condensin, and the SMC5/6 complex in the 59 eukaryotes. For the subunits of the SMC5/6 complex, only Nse1-4 were searched for in this study, because the amino acid sequences of Nse5 and Nse6 were little conserved amongst eukaryotes (Diaz & Pecinka, 2018). The BLAST searches of the aforementioned proteins were conducted as described above. The human homologs are used as the queries in the surveys described above; Rad21/SCC1 (GenBank accession number O60216), STAG1/SCC3 (Q8WVM7), CAP-H (Q15003), CAP-D2 (Q15021), CAP-G (Q9BPX3), Nse1 (Q8WV22), Nse2 (Q96MF7), Nse3 (Q96MG7), and Nse4 (Q9NXX6).

We aligned 126 amino acid sequences belonging to the kleisin superfamily (i.e., 39 Rad21/Scc1, 33 CAP-H, 26 Nse4, 12 archaeal ScpA, and 16 bacterial ScpA sequences) as described above. After the exclusion of ambiguously aligned positions (see above), the alignment was subjected to the ML analysis with the LG + Γ + I model. UFBP support values were calculated from 1,000 replicates.

## ACKNOWLEDGEMENTS

We thank Dr. T. Nakayama (University of Tsukuba, Japan) for his comments and suggestions to the manuscript. This work was supported by the grants from the Japanese Society for Promotion of Sciences awarded to Y. I. (numbers 18KK0203 and 19H03280).

## Data Availability Statement

The alignments used for the phylogenetic analyses, accession numbers of the sequences included in the alignments, and the treefiles from the phylogenetic analyses are available at https://orcid.org/0000-0003-0101-8483.

## REFERENCE

Anderson DE, Losada A, Erickson HP, Hirano T. 2002. Condensin and cohesin display different arm conformations with characteristic hinge angles. J. Cell Biol. 156(3):419–424.

Andrews EA, Palecek J, Sergeant J, Taylor E, Lehmann AR, Watts FZ. 2005. Nse2, a component of the Smc5-6 Complex, Is a SUMO ligase required for the response to DNA damage. Mol. Cell. Biol. 25(1):185–196.

Aragón L. 2018. The Smc5/6 complex: New and old functions of the enigmatic long-distance relative. Annu. Rev. Genet. 52(1):89–107.

Bergsten J. 2005. A review of long-branch attraction. Wiley Online Library.

Birkenbihl RP, Subramani S. 1995. The rad21 gene product of Schizosaccharomyces pombe is a nuclear, cell cycle-regulated phosphoprotein. J. Biol. Chem. 270(13):7703–7711.

Britton RA, Lin DC, Grossman AD. 1998. Characterization of a prokaryotic SMC protein involved in chromosome partitioning. Genes Dev. 12(9):1254–1259.

Camacho C, Coulouris G, Avagyan V, Ma N, Papadopoulos J, Bealer K, Madden TL. 2009. BLAST+: architecture and applications. BMC Bioinform. 10(1):421.

Capella-Gutierrez S, Silla-Martinez JM, Gabaldon T. 2009. trimAl: a tool for automated alignment trimming in large-scale phylogenetic analyses. Bioinformatics. 25(15):1972–1973.

Carramolino L, Lee B, Zaballos A, Peled A, Barthelemy I, Shav-Tal Y, Prieto I, Carmi P, Gothelf Y, González de Buitrago G, Aracil M, Márquez G, Barbero J, Zipori D. 1997. SA-1, a nuclear protein encoded by one member of a novel gene family: molecular cloning and detection in hemopoietic organs. Gene. 195(2):151–159.

Chen S, Zhou Y, Chen Y, Gu J. 2018. fastp: an ultra-fast all-in-one FASTQ preprocessor. Bioinformatics. 34(17):884–890.

Cobbe N, Heck MM. 2000. Review: SMCs in the world of chromosome biology— from prokaryotes to higher eukaryotes. J. Struct. Biol. 129(2-3):123–143.

Cobbe N, Heck MM. 2004. The evolution of SMC proteins: phylogenetic analysis and structural implications. Mol. Biol. Evol. 21(2):332–347.

Diaz M, Pecinka A. 2018. Scaffolding for repair: understanding molecular functions of the SMC5/6 complex. Genes. 9(1):36.

Fousteri MI, Lehmann AR. 2000. A novel SMC protein complex in Schizosaccharomyces pombe contains the Rad18 DNA repair protein. EMBO J. 19(7):1691–1702.

Fu L, Niu B, Zhu Z, Wu S, Li W. 2012. CD-HIT: accelerated for clustering the next-generation sequencing data. Bioinformatics. 28(23):3150–3152.

Fujioka Y, Kimata Y, Nomaguchi K, Watanabe K, Kohno K. 2002. Identification of a novel non-structural maintenance of chromosomes (SMC) component of the SMC5-SMC6 complex involved in DNA repair. J. Biol. Chem. 277(24):21585–21591.

Funayama T, Narumi I, Kikuchi M, Kitayama S, Watanabe H, Yamamoto K. 1999. Identification and disruption analysis of the recN gene in the extremely radioresistant bacterium Deinococcus radiodurans. Mutat. Res. 435(2):151–161.

Gao F, Warren A, Zhang Q, Gong J, Miao M, Sun P, Xu D, Huang J, Yi Z, Song W. 2016. The All-data-based evolutionary hypothesis of ciliated protists with a revised classification of the phylum Ciliophora (Eukaryota, Alveolata). Sci. Rep. 6(1):24874

Gluenz E, Sharma R, Carrington M, Gull K. 2008. Functional characterization of cohesin subunit SCC1 in Trypanosoma brucei and dissection of mutant phenotypes in two life cycle stages. Mol. Microbiol. 69(3):666–680

Grabherr MG, Haas BJ, Yassour M, Levin JZ, Thompson DA, Amit I, Adiconis X, Fan L, Raychowdhury R, Zeng Q, Chen Z, Mauceli E, Hacohen N, Gnirke A, Rhind N, di Palma F, Birren BW, Nusbaum C, Lindblad-Toh K, Friedman N, Regev A. 2011. Full-length transcriptome assembly from RNA-Seq data without a reference genome. Nat. Biotechnol. 29(7):644–652.

Haering CH, Farcas A, Arumugam P, Metson J, Nasmyth K. 2008. The cohesin ring concatenates sister DNA molecules. Nature. 454(7202):297–301.

Hirano T, Mitchison TJ. 1994. A heterodimeric coiled-coil protein required for mitotic chromosome condensation in vitro. Cell. 79(3):449–458.

Hirano T, Kobayashi R, Hirano M. 1997. Condensins, chromosome condensation protein complexes containing XCAP-C, XCAP-E and a Xenopus homolog of the drosophila barren protein. Cell. 89(4):511–521.

Hu B, Liao C, Millson SH, Mollapour M, Prodromou C, Pearl LH, Piper PW, Panaretou B. 2005. Qri2/Nse4, a component of the essential Smc5/6 DNA repair complex. Mol. Microbiol. 55(6):1735–1750.

Ishiguro K. 2019. The cohesin complex in mammalian meiosis. Genes Cells. 24(1):6–30.

Janouškovec J, Paskerova GG, Miroliubova TS, Mikhailov KV, Birley T, Aleoshin VV, Simdyanov TG. 2019. Apicomplexan-like parasites are polyphyletic and widely but selectively dependent on cryptic plastid organelles. eLife. 8:e49662

Katoh K, Standley DM. 2013. MAFFT multiple sequence alignment software version 7: improvements in performance and usability. Mol. Biol. Evol. 30(4):772–780.

Lartillot N, Lepage T, Blanquart S. 2009. PhyloBayes 3: a Bayesian software package for phylogenetic reconstruction and molecular dating. Bioinformatics. 25(17):2286–2288.

Lax G, Eglit Y, Eme L, Bertrand EM, Roger AJ, Simpson AG. 2018. Hemimastigophora is a novel supra-kingdom-level lineage of eukaryotes. Nature. 564(7736):410–414.

Lehmann AR, Walicka M, Griffiths DJ, Murray JM, Watts FZ, McCready S, Carr AM. 1995. The rad18 gene of Schizosaccharomyces pombe defines a new subgroup of the SMC superfamily involved in DNA repair. Mol. Cell. Biol. 15(12):7067–7080.

Losada A, Hirano M, Hirano T. 1998. Identification of Xenopus SMC protein complexes required for sister chromatid cohesion. Genes Dev. 12(13):1986–1997.

Löwe J, Cordell SC, van den Ent F. 2001. Crystal structure of the SMC head domain: an ABC ATPase with 900 residues antiparallel coiled-coil. J. Mol. Biol. 306(1):25–35.

Melby TE, Ciampaglio CN, Briscoe G, Erickson HP. 1998. The symmetrical structure of structural maintenance of chromosomes (SMC) and MukB proteins: long, antiparallel coiled coils, folded at a flexible hinge. J. Cell Biol. 142(6):1595–1604.

Nguyen L, Schmidt HA, von Haeseler A, Minh BQ. 2015. IQ-TREE: a fast and effective stochastic algorithm for estimating maximum-likelihood phylogenies. Mol. Biol. Evol. 32(1):268–274.

Niki H, Jaffé A, Imamura R, Ogura T, Hiraga S. 1991. The new gene mukB codes for a 177 kd protein with coiled-coil domains involved in chromosome partitioning of E. coli. EMBO J. 10(1):183–193.

Palecek JJ, Gruber S. 2015. Kite proteins: A superfamily of SMC/Kleisin partners conserved across Bacteria, Archaea, and Eukaryotes. Structure. 1;23(12):2183–2190.

Palou R, Dhanaraman T, Marrakchi R, Pascariu M, Tyers M, D’Amours D. 2018. Condensin ATPase motifs contribute differentially to the maintenance of chromosome morphology and genome stability. PLOS Biol. 16(6):e2003980.

Pebernard S, McDonald WH, Pavlova Y, Yates JR, Boddy MN. 2004. Nse1, Nse2, and a novel subunit of the Smc5-Smc6 complex, Nse3, play a crucial role in meiosis. Mol. Biol. Cell. 15(11):4866–4876.

Pebernard S, Wohlschlegel J, McDonald WH, Yates 3rd JR, Boddy MN. 2006. The Nse5-Nse6 dimer mediates DNA repair roles of the Smc5-Smc6 complex. Mol Cell Biol. 26(5):1617–30.

Sarai C, Tanifuji G, Nakayama T, Kamikawa R, Takahashi K, Yazaki E, Matsuo E, Miyashita H, Ishida K, Iwataki M, Inagaki Y. 2020. Dinoflagellates with relic endosymbiont nuclei as models for elucidating organellogenesis. Proc. Natl. Acad. Sci. U.S.A. 117(10):5364–5375.

Sayers EW, Beck J, Brister JR, Bolton EE, Canese K, Comeau DC, Funk K, Ketter A, Kim S, Kimchi A, Kitts PA, Kuznetsov A, Lathrop S, Lu Z, McGarvey K, Madden TL, Murphy TD, O’Leary N, Phan L, Schneider VA, Thibaud-Nissen F, Trawick BW, Pruitt KD, Ostell J. 2020. Database resources of the National Center for Biotechnology Information. Nucleic Acids Res. 48(D1):D9–D16.

Schleiffer A, Kaitna S, Maurer-Stroh S, Glotzer M, Nasmyth K, Eisenhaber F. 2003. Kleisins: a superfamily of bacterial and eukaryotic SMC protein partners. Mol Cell. 11(3):571–5.

Shimodaira H. 2002. An approximately unbiased test of phylogenetic tree selection. Syst Biol. 51(3):492–508

Soppa J. 2001. Prokaryotic structural maintenance of chromosomes (SMC) proteins: distribution, phylogeny, and comparison with MukBs and additional prokaryotic and eukaryotic coiled-coil proteins. Gene. 278(1-2):253–264.

Susko E, Roger AJ. 2007. On reduced amino acid alphabets for phylogenetic inference. Mol Biol Evol. 24(9):2139–50.

Sutani T, Yanagida M. 1997. DNA renaturation activity of the SMC complex implicated in chromosome condensation. Nature. 388(6644):798–801.

Takemata N, Samson RY, Bell SD. 2019. Physical and functional compartmentalization of archaeal chromosomes. Cell. 179(1):165–179.

T□th A, Ciosk R, Uhlmann F, Galova M, Schleiffer A, Nasmyth K. 1999. Yeast cohesin complex requires a conserved protein, Eco1p(Ctf7), to establish cohesion between sister chromatids during DNA replication. Genes Dev. 13(3):320–333.

Williams TA, Cox CJ, Foster PG, Szöllősi GJ, Embley TM. 2020. Phylogenomics provides robust support for a two-domains tree of life. Nat. Ecol. Evol. 4(1):138–147.

Yazaki E, Ishikawa SA, Kume K, Kumagai A, Kamaishi T, Tanifuji G, Hashimoto T, Inagaki Y. 2017. Global Kinetoplastea phylogeny inferred from a large-scale multigene alignment including parasitic species for better understanding transitions from a free-living to a parasitic lifestyle. Genes Genet. Syst. 92(1):35–42.

Zaremba-Niedzwiedzka K, Caceres EF, Saw JH, Bäckström D, Juzokaite L, Vancaester E, Seitz KW, Anantharaman K, Starnawski P, Kjeldsen KU, Stott MB, Nunoura T, Banfield JF, Schramm A, Baker BJ, Spang A, Ettema TJ. 2017. Asgard archaea illuminate the origin of eukaryotic cellular complexity. Nature. 541(7637):353–358.

