## Supplementary figures 1-7 for "Ubiquity and origins of structural maintenance of chromosomes (SMC) proteins in eukaryotes"

### ABSTRACT

Structural maintenance of chromosomes (SMC) protein complexes are common in Bacteria, Archaea, and Eukaryota. SMC proteins, together with the proteins related to SMC (SMC-related proteins), constitute a superfamily of ATPases. Bacteria/Archaea and Eukaryotes are distinctive from one another in terms of the repertory of SMC proteins. A single type of SMC protein is dimerized in the bacterial and archaeal complexes, whereas eukaryotes possess six distinct SMC subfamilies (SMC1-6), constituting three heterodimeric complexes, namely cohesin, condensin, and SMC5/6 complex. Thus, to bridge the homodimeric SMC complexes in Bacteria and Archaea to the heterodimeric SMC complexes in Eukaryota, we need to invoke multiple duplications of an SMC gene followed by functional divergence. However, to our knowledge, the evolution of the SMC proteins in Eukaryota had not been examined for more than a decade. In this study, we reexamined the ubiquity of SMC1-6 in phylogenetically diverse eukaryotes that cover the major eukaryotic taxonomic groups recognized to date and provide two novel insights into the SMC evolution in eukaryotes. First, multiple secondary losses of SMC5 and SMC6 occurred in the eukaryotic evolution. Second, the SMC proteins constituting cohesin and condensin (i.e., SMC1-4), and SMC5 and SMC6 were derived from closely related but distinct ancestral proteins. Based on the above-mentioned findings, we discuss how SMC1-6 have diverged from the archaeal homologs.

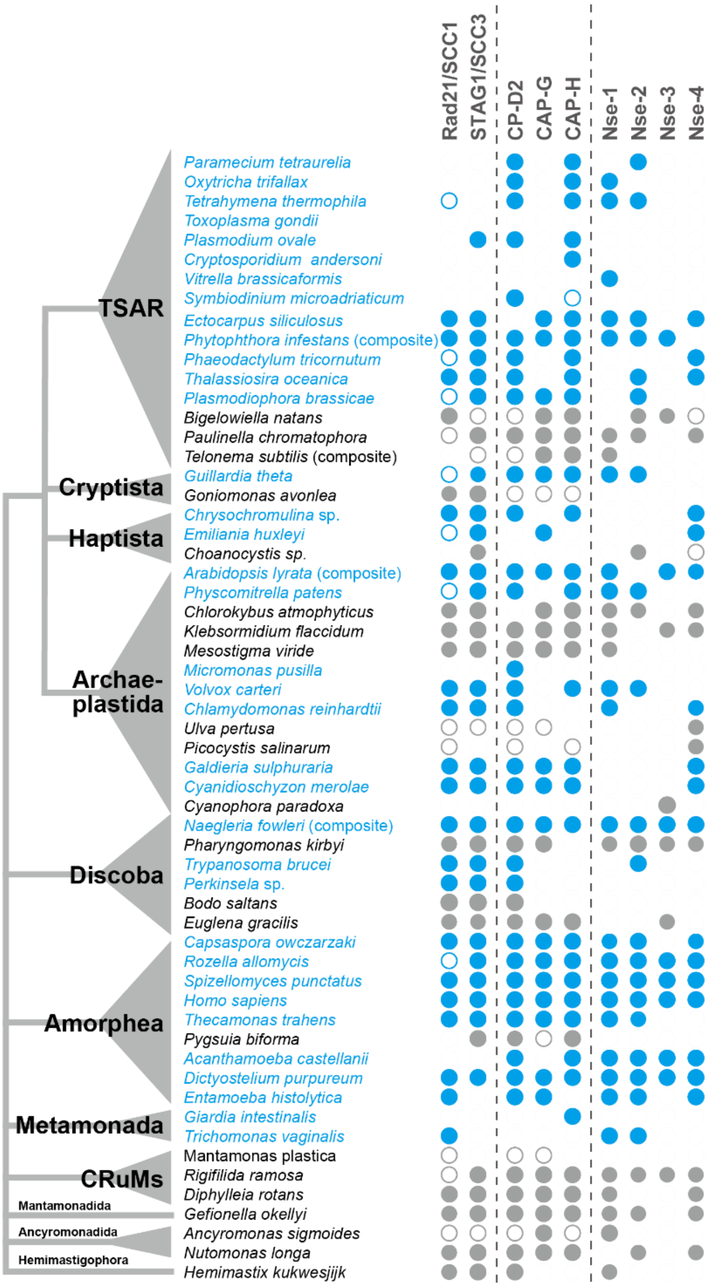


**Supplementary Figure 1.** Inventories of the accessory subunits in the SMC complexes in 59 eukaryotes. The subunits surveyed in this study are Rad21 and Scc3 in cohesin, CAP-D2, CAP-H, and CAP-G in condensin, and Nse1, Nse2, Nse3, and Nse4 in the SMC5/6 complex. Open circles with blue borders and those with grey borders represent the partial sequences found in the GenBank nr database and individual transcriptome data, respectively. Blue and grey (filled) circles represent the putative full-length sequences found in the GenBank nr database and the corresponding transcriptome data, respectively. Note that the amino acid sequences of Nse1-4 are not conserved well among eukaryotes and we could not judge whether the identified sequences were full-length with confidence. Thus, we put light-blue and light-grey circles when the Nse1-4 sequences were found in the GenBank nr database and individual transcriptome data, respectively.

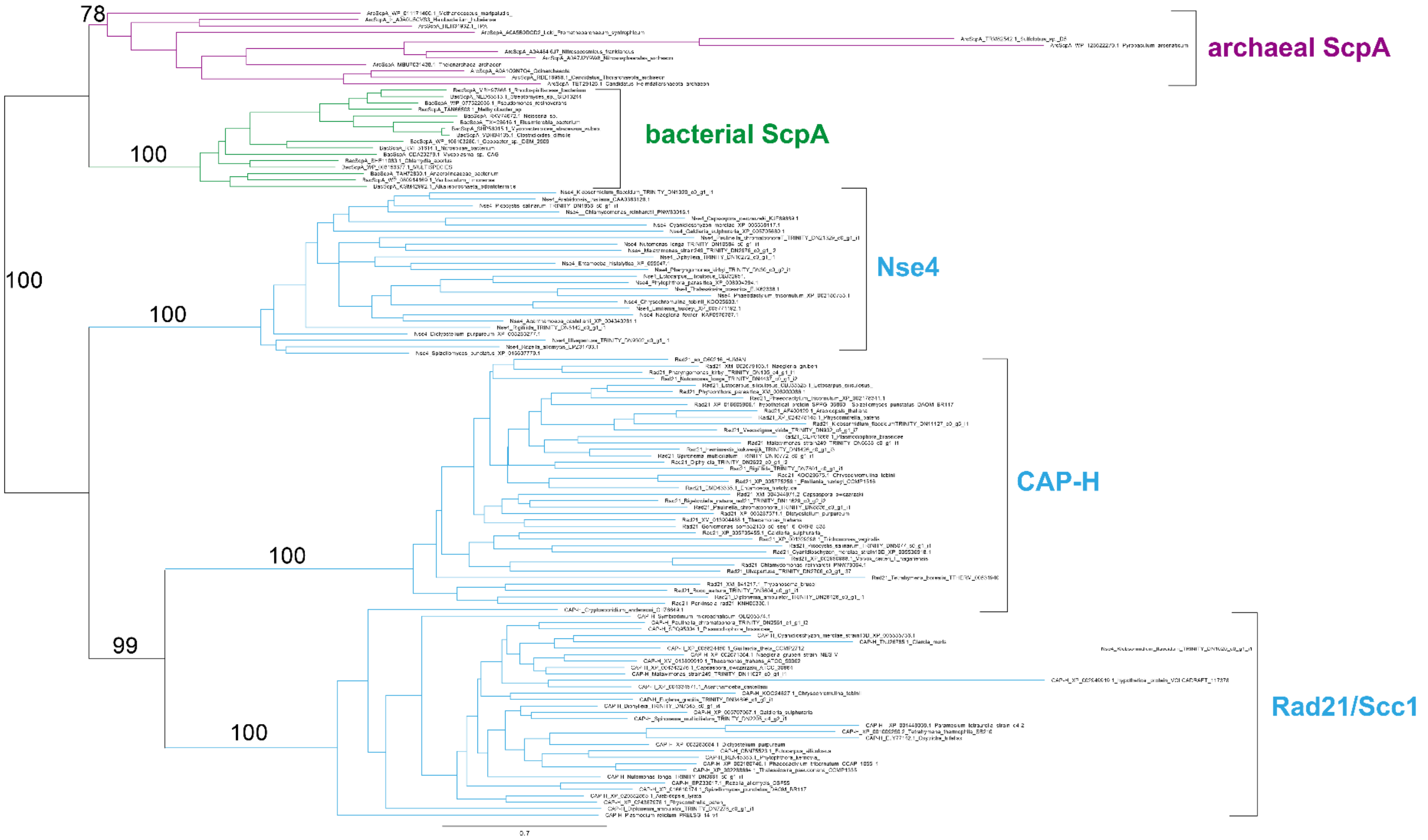


**Supplementary Figure 2.** Maximum-likelihood phylogeny of the kleisin superfamily. The eukaryotic branches (i.e., Rad21/Scc1 in cohesin, CAP-H in condensin, and Nse4 in the SMC5/6 complex) are colored in blue. Bacterial and archaeal Scp1 branches are colored in green and purple, respectively. Only for the major nodes, the ultrafast bootstrap support values are presented.

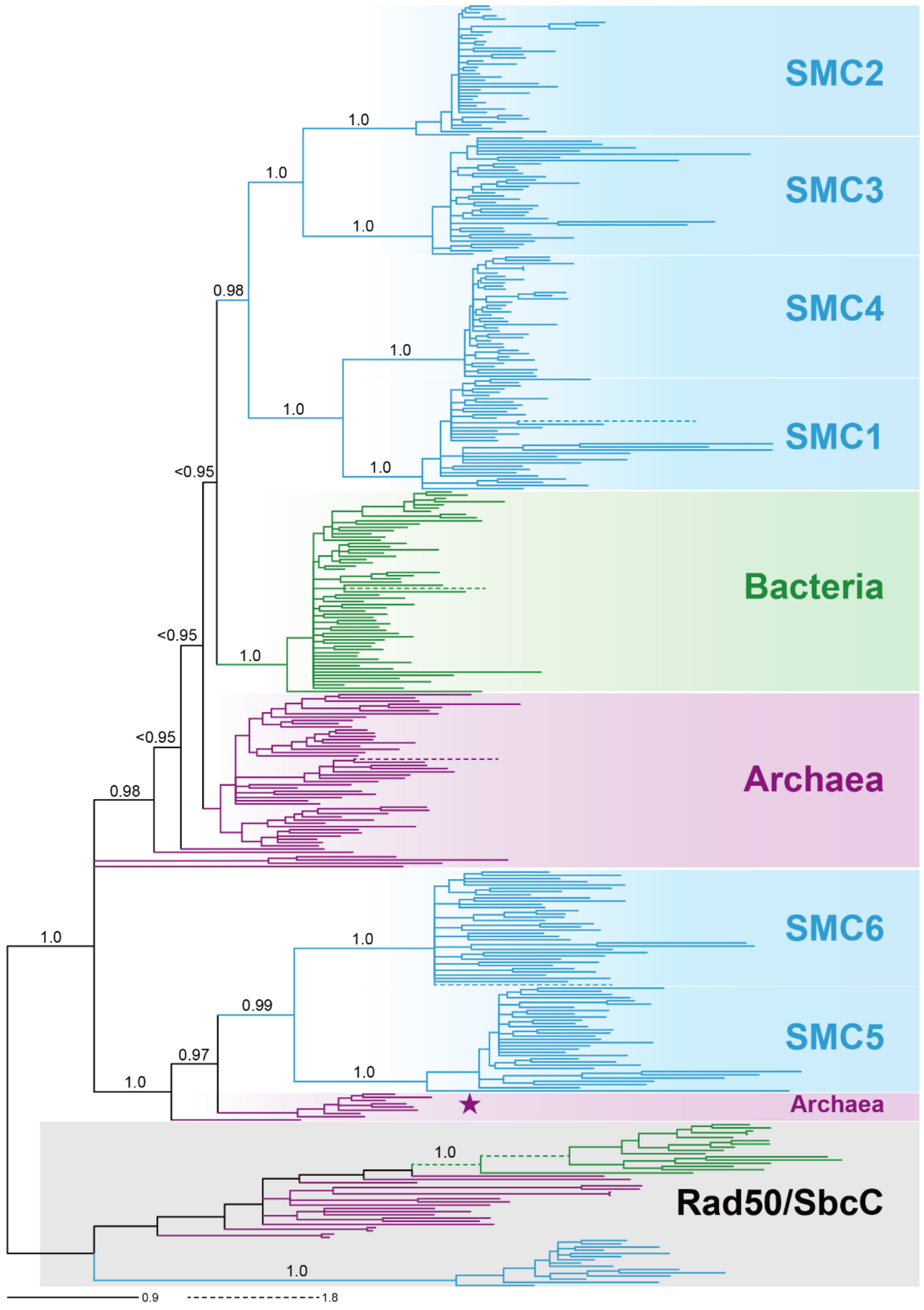


**Supplementary Figure 3.** Bayesian phylogeny of SMC and Rad50/SbcC sequences. All of the sequence names are omitted. The bacterial, archaeal, and eukaryotic branches are colored in green, purple, and blue, respectively. The clade of Rad50/SbcC sequences is shaded in grey. Bayesian posterior probabilities are displayed for the nodes that are critical to the evolution of the six SMC subfamilies of eukaryotes. The 9 archaeal branches are highlighted by a star as “SMC5/6-related archaeal SMC homologs” (see the main text for the details).

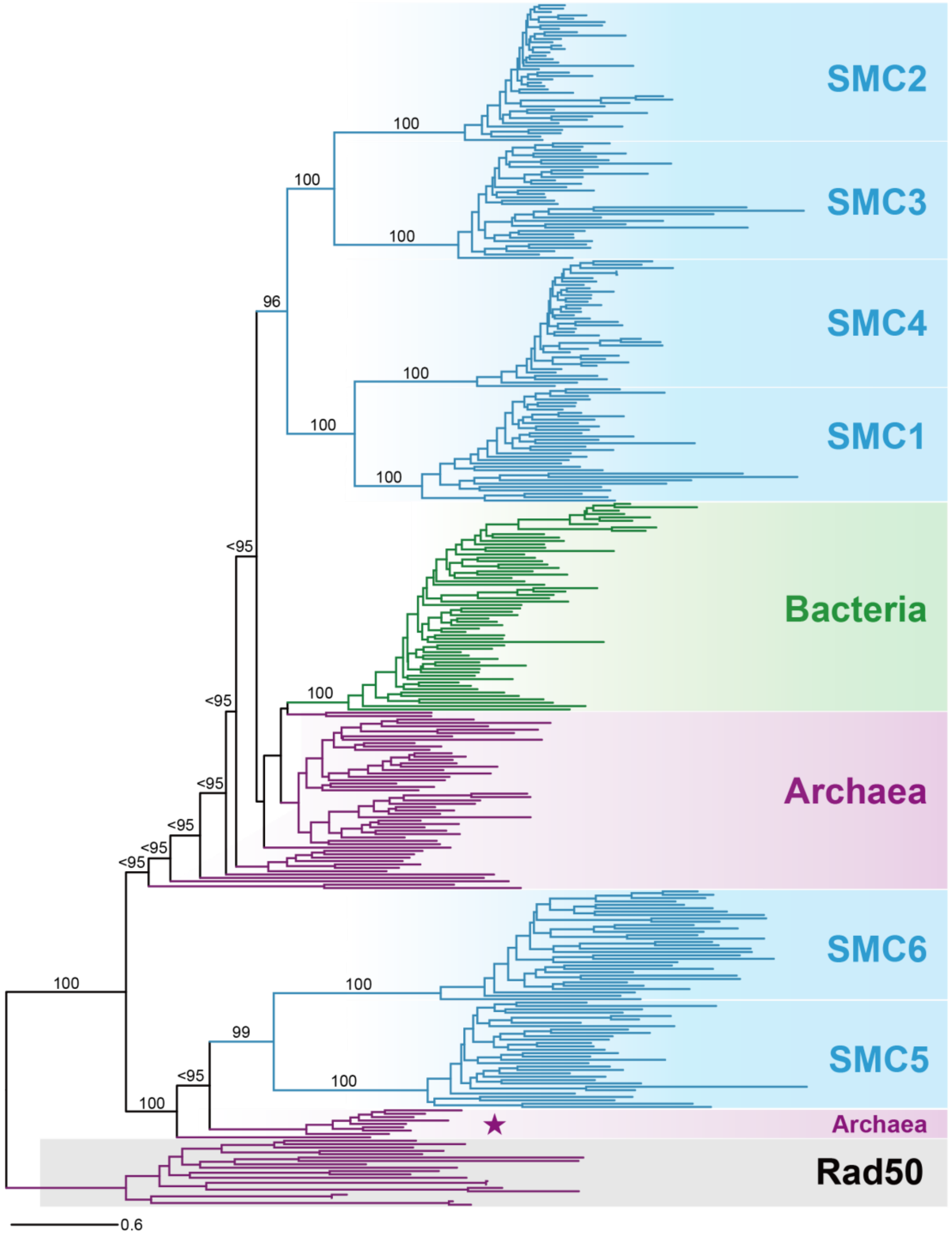


**Supplementary Figure 4.** Maximum likelihood phylogeny of the SMC and Rad50/SbcC sequences after the exclusion of extremely long-branches. All of the sequence names are omitted. The bacterial, archaeal, and eukaryotic branches are colored in green, purple, and blue, respectively. The clade of Rad50 sequences is shaded in grey. Ultrafast bootstrap support values (UFBPs) are displayed for the nodes that are critical to the evolution of the six SMC subfamilies of eukaryotes. Many of the deep splits received UFBPs smaller than 95% (labeled as “<95”). The 9 archaeal branches are highlighted by a star as “SMC5/6-related archaeal SMC homologs” (see the main text for the details).

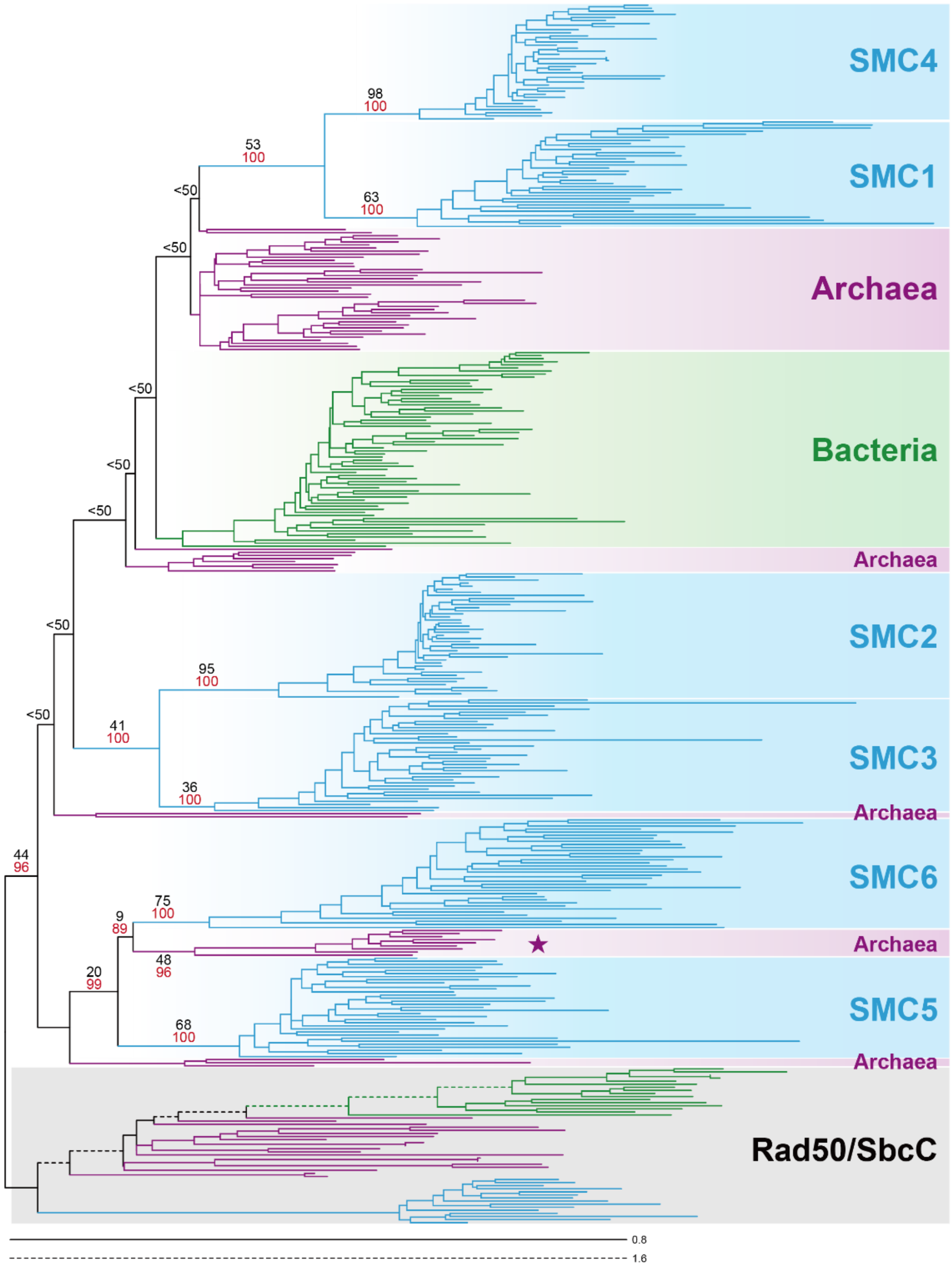


**Supplementary Figure 5.** Maximum likelihood phylogeny of the recoded SMC and Rad50/SbcC sequences. See the main text for the details of the recoding procedure. All of the sequence names are omitted. The bacterial, archaeal, and eukaryotic branches are colored in green, purple, and blue, respectively. The clade of Rad50/SbcC sequences is shaded in grey. Maximum likelihood nonparametric bootstrap support values (MLBPs; colored in black) and ultrafast bootstrap support values (UFBPs; colored in red) are displayed for the nodes that are critical to the evolution of the six SMC subfamilies of eukaryotes. Many of the deep splits received MLBPs smaller than 50% (labeled as “<50”). The 9 archaeal branches are highlighted by a star as “SMC5/6-related archaeal SMC homologs” (see the main text for the details).

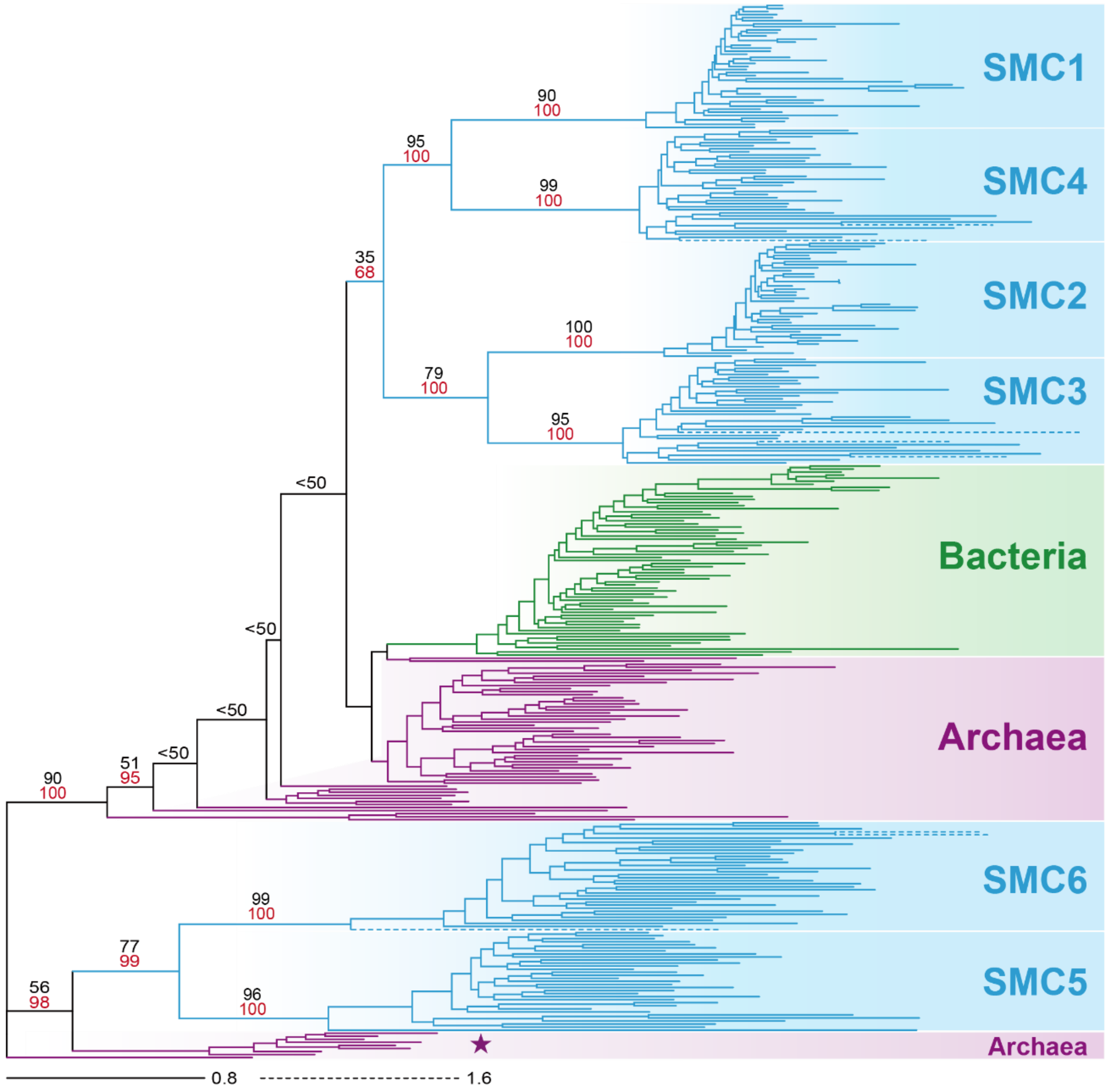


**Supplementary Figure 6.** Maximum likelihood phylogeny inferred from the alignment of the SMC sequences. All of the sequence names are omitted. The bacterial, archaeal, and eukaryotic branches are colored in green, purple, and blue, respectively. Maximum likelihood nonparametric bootstrap support values (MLBPs; colored in black) and ultrafast bootstrap support values (UFBPs; colored in red) are displayed for the nodes that are critical to the evolution of the six SMC subfamilies of eukaryotes. Many of the deep splits received MLBPs smaller than 50% (labeled as “<50”). The 9 archaeal branches are highlighted by a star as “SMC5/6-related archaeal SMC homologs” (see the main text for the details).

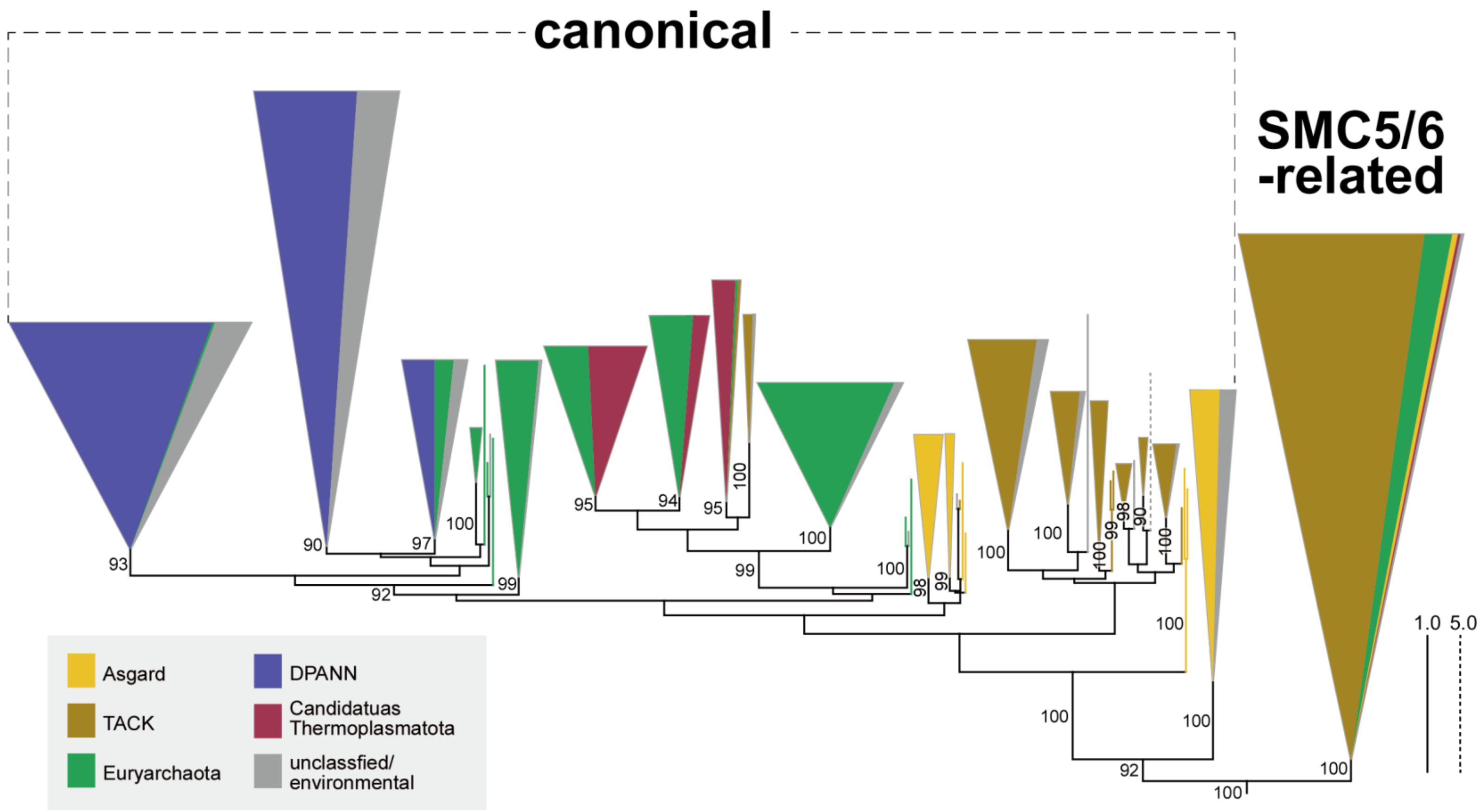


**Supplementary Figure 7.** Maximum likelihood phylogeny of 512 archaeal SMC and 20 Rad50 sequences. All of the sequence names the clade of Rad50 sequences are omitted from this figure. The clades supported by ultrafast bootstrap support values (UFBPs) of ≥90% are collapsed into triangles. Clades and terminal branches are colored according to the Archaeal taxonomy (see the inset for the color-coding). Triangles with multiple colors indicate that the clades comprise the SMC sequences originated form the species/metagenomes belonging to two or more major lineages of Archaea. The sizes of individual colored areas are proportional to the numbers of the sequences of the corresponding lineages in the clade. Only UFBPs ≥90%are indicated.
